# Mapping the vocal circuitry of Alston’s singing mouse with pseudorabies virus

**DOI:** 10.1101/2021.07.16.452718

**Authors:** Da-Jiang Zheng, Daniel E. Okobi, Ryan Shu, Rania Agrawal, Samantha K. Smith, Michael A. Long, Steven M. Phelps

## Abstract

Vocalizations, like many social displays, are often elaborate, rhythmically structured behaviors that are modulated by a complex combination of cues. Vocal motor patterns require close coordination of neural circuits governing the muscles of the larynx, jaw, and respiratory system. In the elaborate vocalization of Alston’s singing mouse (*Scotinomys teguina*), for example, each note of its rapid, frequency-modulated trill is accompanied by equally rapid modulation of breath and gape. To elucidate the neural circuitry underlying this behavior, we introduced the polysynaptic retrograde neuronal tracer pseudorabies virus (PRV) into the cricothyroid and digastricus muscles, which control frequency modulation and jaw opening respectively. Each virus singly labels ipsilateral motoneurons (nucleus ambiguous for cricothyroid, and motor trigeminal nucleus for digastricus). We find that the two isogenic viruses heavily and bilaterally co-label neurons in the gigantocellular reticular formation, a putative central pattern generator. The viruses also show strong co-labeling in compartments of the midbrain including the ventrolateral periaqueductal grey and the parabrachial nucleus, two structures strongly implicated in vocalizations. In the forebrain, regions important to social cognition and energy balance both exhibit extensive co-labeling. This includes the paraventricular and arcuate nuclei of the hypothalamus, the lateral hypothalamus, preoptic area, extended amygdala, central amygdala, and the bed nucleus of the stria terminalis. Finally, we find doubly labeled neurons in M1 motor cortex previously described as laryngeal, as well as in the prelimbic cortex, which indicate these cortical regions play a role in vocal production. Although we observe some novel patterns of double-labelling, the progress of both viruses is broadly consistent with vertebrate-general patterns of vocal circuitry, as well as with circuit models derived from primate literature.

## Introduction

Animal displays are among the most diverse and dramatic behaviors in the natural world, and their roles in courtship, aggression, and other contexts have long been the subject of behavioral studies (Cummings & Endler, 2018). Displays also make excellent foci for neurobiological research (Fusani, Barske, Day, Fuxjager, & Schlinger, 2014; Remage-Healey, Maidment, & Schlinger, 2008). Their motor patterns are complex and require the precise coordination of many muscles, but they are often highly stereotyped. Moreover, tuning a display to its ecological context requires not only the integrated control of muscles, but also the modulation of display by hormonal states and complex social cues (Schlinger, Paul, & Monks, 2018). Identifying the neural circuits that coordinate displays can enable insights into the diversity of complex behaviors and their decision mechanisms.

Among the many forms of display, perhaps none is more widespread or well-studied than vocalization (Barkan & Zornik, 2020; Zhang & Ghazanfar, 2020). Phylogenetic studies reveal that vocalization is common and arose only a few times during vertebrate evolution (Chen & Wiens, 2020). Vertebrate vocalizations function in courtship, aggression, individual identification, parental care and many other contexts (Suthers, Fitch, Fay, & Popper, 2004). Adaptive vocalization is often sensitive to internal states like reproductive status or body condition, as well as to external cues like the immediate social environment (Yamaguchi and Kelley 2003, Gentner & Margoliash, 2006).

Classic studies on vocal mechanisms in primates have combined tract-tracing, electrophysiology, pharmacological manipulation and other methods to identify substrates of vocal behavior, from limbic cortex to the spinal cord (Jurgens 2009). A smaller body of work builds and extends this into model rodent species, such as the lab mouse and lab rat (Bennett, Maier, & Brecht, 2019; Tschida et al., 2019). Overall, this literature suggests some surprising commonalities to rodent and primate vocal circuitry, but too few species have been sampled to enable robust inferences about general mammalian patterns. Among tetrapods more generally, work has often focused on circuits motivated by research traditions within specific study taxa. Among songbirds, for example, the emphasis has often been on forebrain circuits for song-learning (Brainard & Doupe, 2013) and production (Fee et al., 2004). In African clawed frogs and in the plainfin midshipman fish, vocal work has focused on motor mechanisms at the level of the brainstem and lower, and on their coordination by limbic structures in the hypothalamus and amygdala (Goodson & Bass, 2002; Hall, Ballagh, & Kelley, 2013). Although the research is often piecemeal, together the results suggest features of neural organization that may be shared across vocal vertebrates. Here we seek to more exhaustively elucidate the vocal circuitry in a novel mammalian model, Alston’s singing mouse (*Scotinomys teguina*), by performing viral tract-tracing targeting specific vocal muscles. We compare these findings to reports in mammals and other tetrapods.

Alston’s singing mice are small muroid rodents in the family Cricetidae, which make long, relatively loud vocalizations in the range of human hearing (Campbell et al., 2010; Hooper & Carleton, 1976; Miller & Engstrom, 2007). During vocalization, animals assume an upright posture, open their mouths, and angle their snouts upward (Banerjee, Phelps, & Long, 2019; Hooper & Carleton, 1976; Riede & Pasch, 2020). Each song consists of a series of frequency-modulated notes (Campbell et al., 2010). Each note requires opening the mouth and making a short exhalation that accompanies vocalization (Okobi, Banerjee, Matheson, Phelps, & Long, 2019; Pasch, George, Campbell, & Phelps, 2011a). The vocalization is highly stereotyped, with each song consisting of frequency-modulated notes that lengthen regularly as the song progresses (Campbell et al., 2010). The song is used for female attraction (Fernández-Vargas, Tang-Martínez, & Phelps, 2011), male-male aggression (Pasch, George, Campbell, & Phelps, 2011b), and species recognition (Pasch, Bolker, & Phelps, 2013). It is modulated by reproductive state, stress reactivity and energy balance (Burkhard, Westwick, & Phelps, 2018; Crino, Larkin, & Phelps, 2010; Pasch et al., 2011a, Giglio and Phelps 2020). Like many displays, singing would seem to require a network of brain regions that integrate diverse cues and translate them into adaptive, rhythmic motor patterns. Although the singing mouse is a compelling species for behavioral study, we know relatively little about the neural mechanisms of *Scotinomys* vocalization.

To quickly elucidate the vocal circuits of *Scotinomys teguina,* we employed the pseudorabies virus bartha (PRV-Bartha; Card & Enquist, 2014). PRV-Bartha is a strain of alpha-herpes virus that has been attenuated for the use of retrograde trans-synaptic transport in mammalian neurons (Pickard et al 2002, Card and Enquist 2014). This trans-synaptic spread enables us to examine multiple candidate brain structures. In addition, the time of arrival of the virus in a specific region allows a preliminary estimation of the network topology (Banfield et al 2003). Lastly, the availability of multiple isogenic strains of PRV-Bartha allows dual labeling of neurons that shape activity of different end-organs (Hogue, Card, Rinaman, Staniszewska Goraczniak, & Enquist, 2018). Although the dual isogenic PRV approach has been used extensively to examine the control of feeding, homeostasis and sympathetic function (Doslikova et al., 2019; Pérez et al., 2011; Stanley et al., 2010; Wee, Lorenz, Bekirov, Jacquin, & Scheller, 2019; Wiedmann, Stefanidis, & Oldfield, 2017), to our knowledge it has not been used in the context of vocalization.

In the current study, we use a dual-label approach to infect two distinct muscles that play essential roles in the vocalizations of singing mice. The first target is the cricothyroid muscle, an intrinsic muscle of the larynx important to frequency modulation (Riede, 2013). The second target is the digastricus muscle, which opens the jaw. Because these two muscles must be coordinated to produce the regularly repeated notes of a *Scotinomys* song, we reasoned that identifying neurons that were double-labelled would reveal circuits involved in vocalization. To do so, we injected one PRV-Bartha strain expressing GFP, and another expressing RFP into either the jaw or the larynx. The two injections were ipsilateral and of equal viral titers. We describe how viruses injected into each muscle arrive into regions of the vocal circuit over time, and we quantify the number of cells each virus infects as well as the number of co-infected cells. Finally, we compare the order of arrival of these viruses to known network topologies in other species. Together these data allow us to quickly characterize the vocal circuitry of a novel model for mammalian vocalization, Alston’s singing mouse.

## Methods

### Animals

We used 16 male *S.teguina* outbred from a wild population caught near Quetzal Education Research Center (QERC), in San Gerardo de Dota, Costa Rica. We have reported housing and husbandry previously in Zheng et al (2021). All animal protocols were approved by the IACUC committee at The University of Texas at Austin in accordance with the National Institute of Health *Guide for the Care and Use of Laboratory Animals*. *S.teguina* weighted 15.2 ± 1.5 grams and were at 143 ± 28 days of age at the time of viral inoculation. Animals were housed in a biosafety level 2 facility.

### Pseudorabies virus

We obtained USDA Aphis approval (Permit # 135766) allowing for interstate shipment of PRV Bartha from Princeton University to The University of Texas at Austin and from UT Austin’s Institutional Biosafety Committee to acquire and employ the virus in our facility. We received two isogenic recombinants of the Bartha strain of PRV from the Center for Neuroanatomy and Neurotropic Viruses. PRV-152 expressing eGFP (enhanced green fluorescent protein, titer:2.45*10^9^ pfu/ml) and PRV-614 expressing mRFP1 (monomeric red fluorescent protein, titer:1.55*10^9^ pfu/ml). Upon receipt of the stocks, the virus were aliquoted at 20μl/cryovial in a biosafety cabinet, flash frozen, and stored in a locked −80°C freezer. On the day of surgery, individual cryovials were placed in a dedicated freezing container (Nunc) and thawed immediately before injection. Excess virus was inactivated by 10% bleach at a ratio of 10-to-1 and disposed in biosafety waste.

Subjects were anesthetized using isoflurane (5% induction, 2.5% maintenance) mixed with oxygen at a flow rate of 1ml/min. A dedicated anesthetist monitored and recorded the anesthesia plane during surgery. The animal was placed on a Kopf 900 stereotax on top of an infrared heating pad (Kent Scientific). The stereotax fitted with a 923-B mouse gas anesthesia head holder (Kopf instruments) allowing for rotation along the vertical axis. Once rotated, a small amount of depilatory cream (Nair) was applied in the jaw and throat area for less than one minute. The area was then sterilized with an alternating application of providone-iodine and 70% ethanol. Analgesics including carprofen, slow-releasing buprenorphine, and lidocaine were applied.

Given the difference in titer between the two strains, we adjusted injection volumes to deliver an approximately equal titer of each virus (500nl PRV 154, 700nl PRV-614, ∼1.1-1.2*10^3^ pfu). Viruses were always given ipsilaterally. The virus assigned to each muscle, and the side of the body targeted (left or right) was randomized and counter-balanced.

We first targeted the cricothyorid muscle, which was exposed through a 1cm incision, accessing the sternohyoid muscle. A 0.5cm incision was made to expose the larynx and its internal muscles. Overall, our cricothyroid injection approach was similar to a previously reported surgical procedure (Arriaga, Macopson, & Jarvis, 2015). We used a nanoliter injector (Nanoject III, Drummond) to deliver a predetermined amount of the PRV into the muscle. The injector was fitted with a pulled (P-2000, Sutter Instruments) Wiretrol I calibrated micropipette (Drummond) and forged to a tip of 20μm I.D (Narishige). We injected 100nl boluses at a rate of 3nl per second separated by one minute. After the cricothyroid injection, we would dispose the pipette in bleach and prepare a second pipette for the other PRV strain into the digastricus. Another 1-cm incision was made in the jaw area of the subject, exposing the posterior and anterior digastricus. We injected unilaterally into the anterior digastricus at the same rate as the cricothyroid injection. After delivering the virus, the muscles were dried with a sterile cotton swab and the skin flap was glued together with Vetbond (3M). Animals were housed individually and monitored every 12 hours.

Previous results have demonstrated that double viral injections with these strains have similar infection kinetics as single injections (Banfield, Kaufman, Randall, & Pickard, 2003; Hettigoda et al., 2015; Hogue et al., 2018; Jovanovic, Pastor, & O’Donovan, 2010; Stanley et al., 2010; Wee et al., 2019). In pilot work, we performed unilateral injections targeting the cricothyroid and digastricus muscle separately with the animals sacrificed at 48 and 60 hours post-injection. We observed very little infection in subjects infected for less than 60 hours. Conversely, subjects were moribund at 96 hours post-injection. We first sought to characterize the time-course and extent of infection using six animals at three time points (72h, 84h, & 96h post-infection; n=2 per time point). At 72h, only a few structures in the brainstem and spinal cord were labelled, so we injected a larger group of 10 animals at two time points (84 & 96h post-infection, n=5 per time point).

To quantify patterns of labelling and co-labelling, we selected a subset of brain regions expressing markers for one or both viruses, and spanning a full range of neuroanatomical levels – including regions of the brainstem, midbrain, hypothalamus, amygdala and neocortex. We quantified single- and double-labelling in our final cohort of 96h subjects (n=5), and report qualitative findings for the remainder of our samples. Sample sizes, targeted tissues and virus-muscle assignments are summarized in Table 1.

**Table 1:**
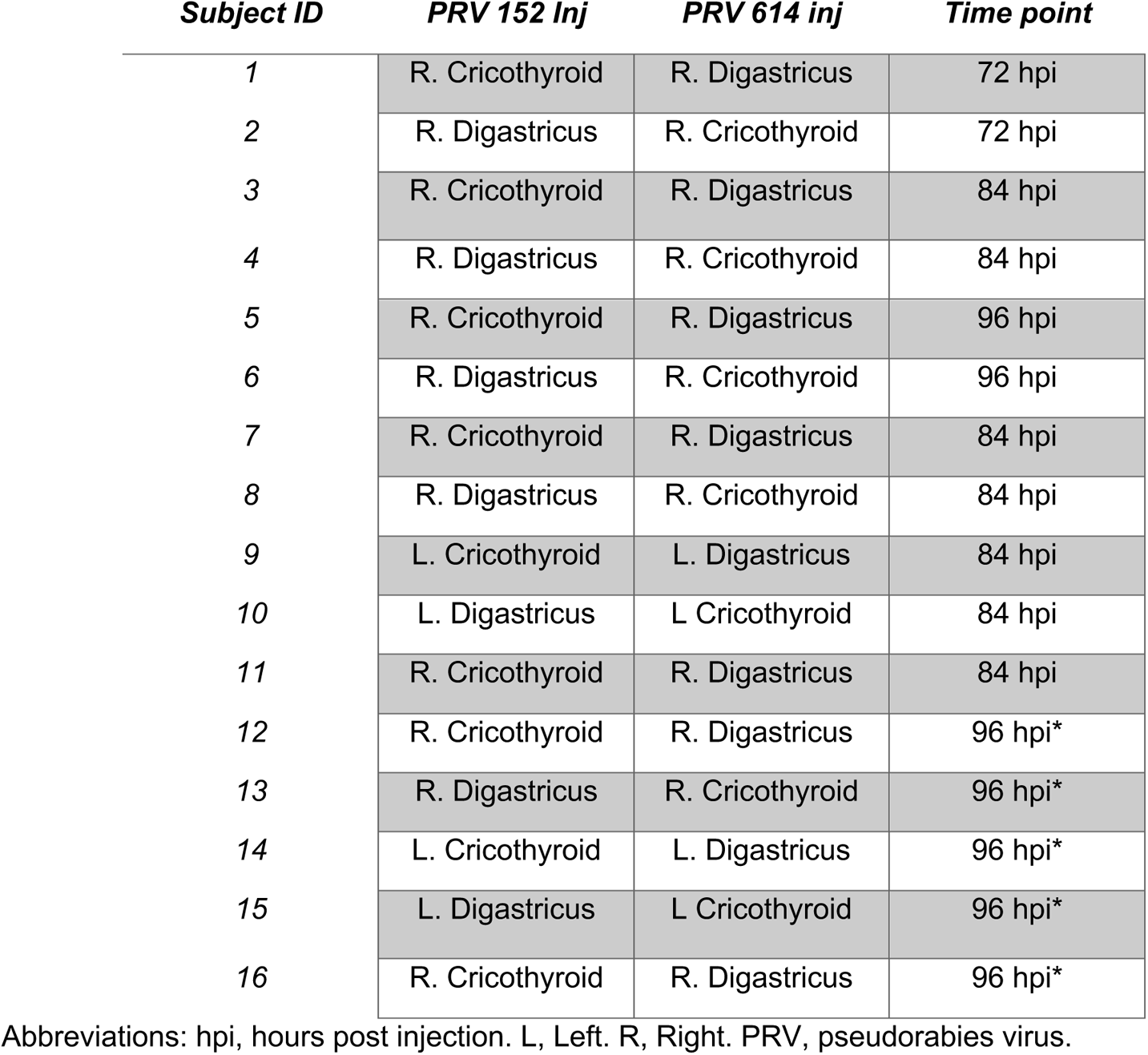
An overiew of PRV type and injection strategy, including survival times included in this study. * denotes subject used in quantitative study.

| ***Subject ID*** | ***PRV 152 Inj*** | ***PRV 614 inj*** | ***Time point*** |
| --- | --- | --- | --- |
| 1 | R. Cricothyroid | R. Digastricus | 72 hpi |
| 2 | R. Digastricus | R. Cricothyroid | 72 hpi |
| 3 | R. Cricothyroid | R. Digastricus | 84 hpi |
| 4 | R. Digastricus | R. Cricothyroid | 84 hpi |
| 5 | R. Cricothyroid | R. Digastricus | 96 hpi |
| 6 | R. Digastricus | R. Cricothyroid | 96 hpi |
| 7 | R. Cricothyroid | R. Digastricus | 84 hpi |
| 8 | R. Digastricus | R. Cricothyroid | 84 hpi |
| 9 | L. Cricothyroid | L. Digastricus | 84 hpi |
| 10 | L. Digastricus | L Cricothyroid | 84 hpi |
| 11 | R. Cricothyroid | R. Digastricus | 84 hpi |
| 12 | R. Cricothyroid | R. Digastricus | 96 hpi* |
| 13 | R. Digastricus | R. Cricothyroid | 96 hpi* |
| 14 | L. Cricothyroid | L. Digastricus | 96 hpi* |
| 15 | L. Digastricus | L Cricothyroid | 96 hpi* |
| 16 | R. Cricothyroid | R. Digastricus | 96 hpi* |

Abbreviations: hpi, hours post injection. L, Left. R, Right. PRV, pseudorabies virus.

### Tissue dissections and cryosectioning

At set time points (Table 1), animals were euthanized by isoflurane to effect, and then transcardially perfused with a peristaltic pump (Cole-Parmer), first with cold 1X PBS (Invitrogen) and then cold 4% paraformaldehyde (EMS) in 1X PBS. Brains were removed, post-fixed in 4% paraformaldehyde for 24 hours at 4°C, and then cryoprotected in 15% followed by 30% sucrose. After cryoprotection, brains were frozen on powdered dry ice and stored in a −80°C freezer until sectioning. We collected alternating series at 30μm with a HM550 cryostat (Thermo-scientific) beginning just caudal to the nucleus ambiguus, corresponding to approximately −7.50mm with reference to Franklin and Paxinos 2007), and extending rostrally until appearance of the accessory olfactory nucleus. For qualitative study, individual sections were placed in a cryoprotectant (de Olmos solution) and stored in −20°C until immunofluorescence. For our quantitative study, individual sections were mounted on a Superfrost Plus slide (Fisher scientific) and stored in −80°C until immunofluorescence. No individual section was exposed to more than one freeze-thaw cycle. We noticed no qualitative difference in brain areas infected between the sections stored in wells and those mounted on slides.

### Immunofluorescence

One of the two series collected was used for multiplex immunofluorescence of eGFP and mRFP signal. All antibodies, ratios, RRIDs, and prior publications using the antibodies are reported in Table 2. In our transition from our timecourse analysis to our focal sample, we changed the RFP antibody to allow us to triple label sections with a specific androgen receptor antibody PG21 (Prins, Birch, & Greene, 1991) – data we have yet to compile, and that are not part of our current study. This antibody change had no discernible effect on our labelling patterns.

**Table 2:** List of antibodies used in this study.

| Target | Host | Dilution | Source | Catalog number | RRID | Citation |
| --- | --- | --- | --- | --- | --- | --- |
| **Primary antibodies** |  |  |  |  |  |  |
| **GFP** | Chicken | 1:1000 | Abcam | Ab13970 | AB_300798 | Schneider et al 2019 |
| **RFP** | Rabbit | 1:500 | Abcam | Ab62341 | AB_945213 | Yao et al 2018 |
| **RFP** | Goat | 1:500 | Rockland | 200-101-379 | AB_2744552 | Chen et al 2017 |
| **Tyrosine hydroxylase** | Sheep | 1:1000 | Millipore | AB1542 | AB_90755 | Greenberg et al 2015 |
| **Secondary antibodies** |  |  |  |  |  |  |
| **Chicken Alexa Fluor 488** | Donkey | 1:200 | Jackson Immuno | 703-545-155 | AB_2340375 |  |
| **Rabbit Alexa Fluor 594** | Donkey | 1:200 | Thermo Fisher | A21207 | AB_141637 |  |
| **Goat Alexa Fluor 594** | Donkey | 1:200 | Thermo Fisher | A11058 | AB_142540 |  |
| **Sheep Alexa Fluor 680** | Donkey | 1:200 | Thermo Fisher | A21102 | AB_1500713 |  |

Abbreviations: GFP, green fluorescent protein. RFP, red fluorescent protein

We labeled the most of the rostral-caudal axis of the *S.teguina* brain. We first washed in 1X PBS for three times and then incubated in a blocking solution with 10% normal serum, 0.3% Triton X, and 1X PBS for one hour. Sections were rinsed again with 1X PBS for three times and then incubated in 5% normal serum, 0.3 Triton X, 1X PBS, with the primary antibody (Table 2) for 24 hours in room temperature. Sections were then rinsed again three times in 1X PBS and then incubated in corresponding secondary antibodies for 2 hours at room temperature (Table 2). Slides were then covered with a hardset with DAPI (Vectashield) and stored in 4°C under dark conditions until microscopy.

### Microscopy and cell counting

All imaging was performed at the Center for Biomedical Research Support at the University of Texas at Austin with a W1 Yokogawa spinning disk confocal microscope (Nikon). Excitation laser power was optimized for 488, 561, and 640nm channels and a Plan Fluor DLL Ph1 10X/0.3NA objective was used to collect all images. Respective laser power was kept consistent through all imaging of tissue samples. For tiling of large images, a scan large-image macro was used with a 20% overlap and blending option. Images collected in this fashion were used for qualitative assessment of infection.

We used both the Paxinos and Franklin third edition atlas (Franklin & Paxinos, 2007) along with the Allen Brain atlas (Lein et al., 2007) to delineate regions of interest. ROIs reported, number of bregma levels, and the estimate bregma level according to Paxinos and Franklin are reported in Table 3. Pseudo-colored images were automatically thresholded for brightness and contrast in ImageJ, and soma were counted using the Cell Counter suite. The merged color was then split by channel and counted again for virus-specific infection. Each image was counted twice, once by the primary scorer (DJZ) and once by a naïve scorer (RA). Muscles were randomly assigned to either the RFP- or GFP containing viruses, and these were counterbalanced across animals. For clarity we present infections by the cricothyroid as green and digastricus as red (MAGENTA) and dual infections as yellow (WHITE).

**Table 3:** List of ROIs quantified in this study.

| ROI | Number of sections counted | Bilateral/Unilateral | Approximate bregma |
| --- | --- | --- | --- |
| **DMPSP5** | 2 | Unilateral | -6.64 |
| **AMB** | 5 | Unilateral | -6.64 |
| **NTS** | 5 | Bilateral | -6.64 |
| **GI** | 5 | Bilateral | -6.64 |
| **LC** | 5 | Bilateral | -5.68 |
| **LPB** | 5 | Bilateral | -5.68 |
| **MPB** | 5 | Bilateral | -5.52 |
| **DMPAG** | 10 | Unilateral | -4.96 |
| **LPAG** | 10 | Bilateral | -4.96 |
| **LH** | 5 | Bilateral | -2.3 |
| **PVN** | 5 | Unilateral | -0.82 |
| **BNST** | 5 | Bilateral | 0.02 |
| **M1** | 5 | Unilateral | -0.10 |
| **PrL** | 3 | Bilateral | 1.94 |

\*Approximate bregma levels are referenced from Franklin and Paxinos (Franklin and Paxinos 2007) from where counting began caudal to rostral in mm.
Abbreviations: AMB, nucleus ambiguus. BNST, bed nucleus of the stria terminalis. DMPSP5, dorsomedial trigeminal nucleus. DMPAG, dorsomedial periaqueductal grey. Gi, gigantocellular reticular nucleus. LC, locus coeruleus. LH, lateral hypothalamus. LPAG, lateral periaqueductal grey. LPB, lateral parabrachial nucleus. M1, primary motor cortex. MPB, medial parabrachial nucleus. NTS, nucleus of the solitary tract. PrL, prelimbic cortex. PVN, paraventricular nucleus. ROI, region of interest.

## Results

The overall goal of our study is to explore the system of interconnected brain regions that innervate two muscles (cricothyroid and digastricus) that are involved in the stereotyped vocalizations of *S.teguina* (Figure 1). The infection by the dual PRVs was both pervasive and specific. Below we describe a systematic survey of label originating from either the cricothyroid, the disgastricus, or both. The description of labelling begins at the caudal boundary of regions that were sampled across individuals. Because there is no atlas of the singing mouse brain, we refer to “bregma levels” with respect to the *Mus musculus* atlas of Franklin and Paxinos (2007) to describe the relative position of the staining. For the subset of structures that we quantified by cell-counting, these levels are also reported in Table 3.

**Figure 1.**
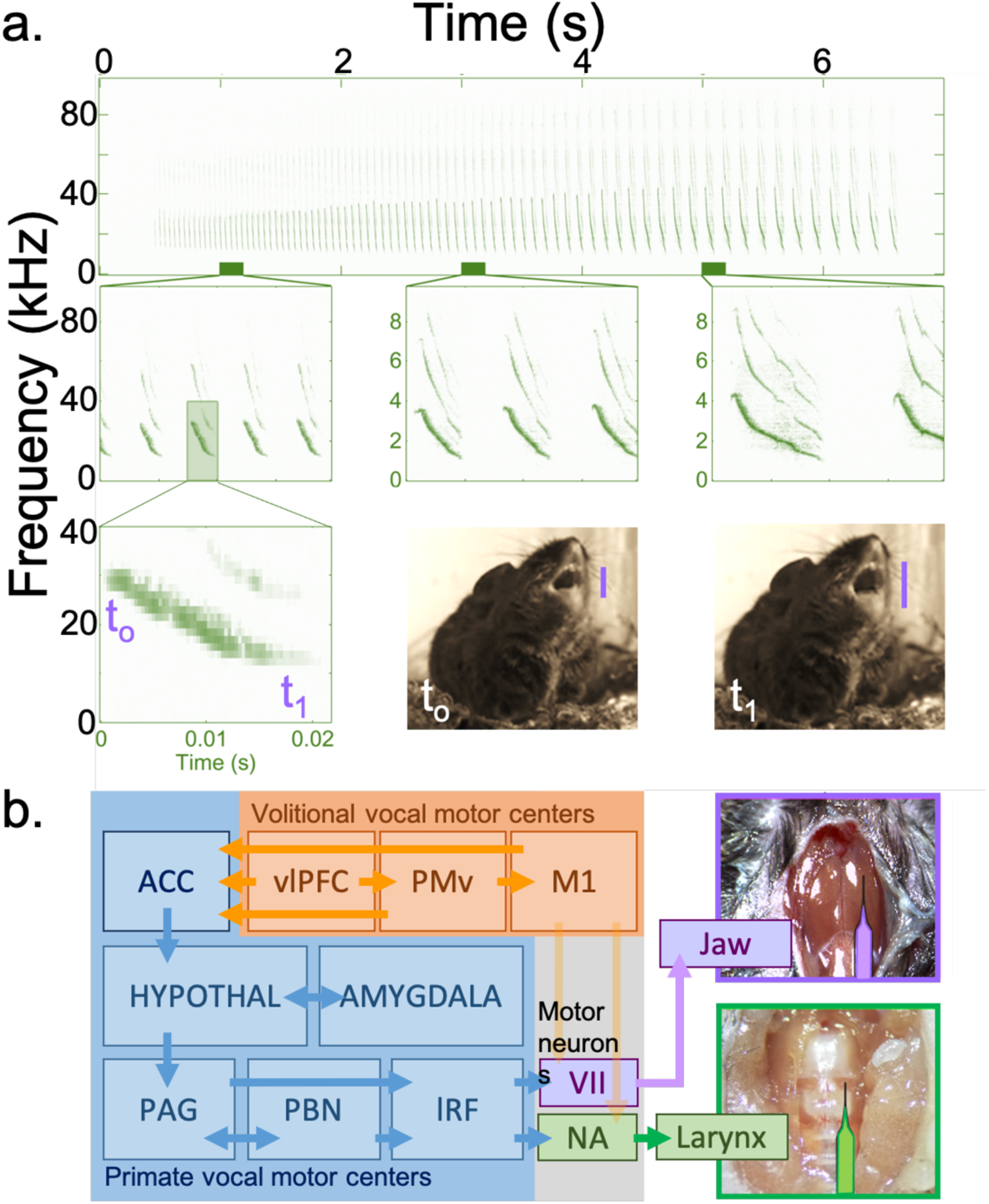
Schematic of experimental approach. 1a, high-speed capture of the advertisement call behavior of *S.teguina*. Vocal gape indicated by vertical lines. Images courtesy of Dr Bret Pasch, NAU. 1b, relative target positions of the muscles in this study. 1c, adaptation of schematic from Hage and Nieder 2016 as applied to this current experiment.

### Brainstem and spinal cord

Cricothyroid injection resulted in a single-labelled, ipsilateral infection in the nucleus ambiguus (Figure 2a). The nucleus was labelled at 72h, the earliest timepoint observed for any neurons labelled with virus injected into the cricothyroid (Figure 2b, c). The infection was restricted to the rostral portion of the nucleus ambiguus, roughly coinciding with the presence of the inferior olive, which is devoid of infection. In contrast, areas surrounding the nucleus ambiguus, particularly the pre-botzinger complex, were labelled by the digrasticus virus at 72h, and co-labelled by both at 84 and 96h (Figure 2a, c). By 96h, a few double-labelled cells were present in the ipsilateral nucleus ambiguous, as well as a small number of labelled contralateral neurons (see Figure 2a,c and Table 4).

**Figure 2.**
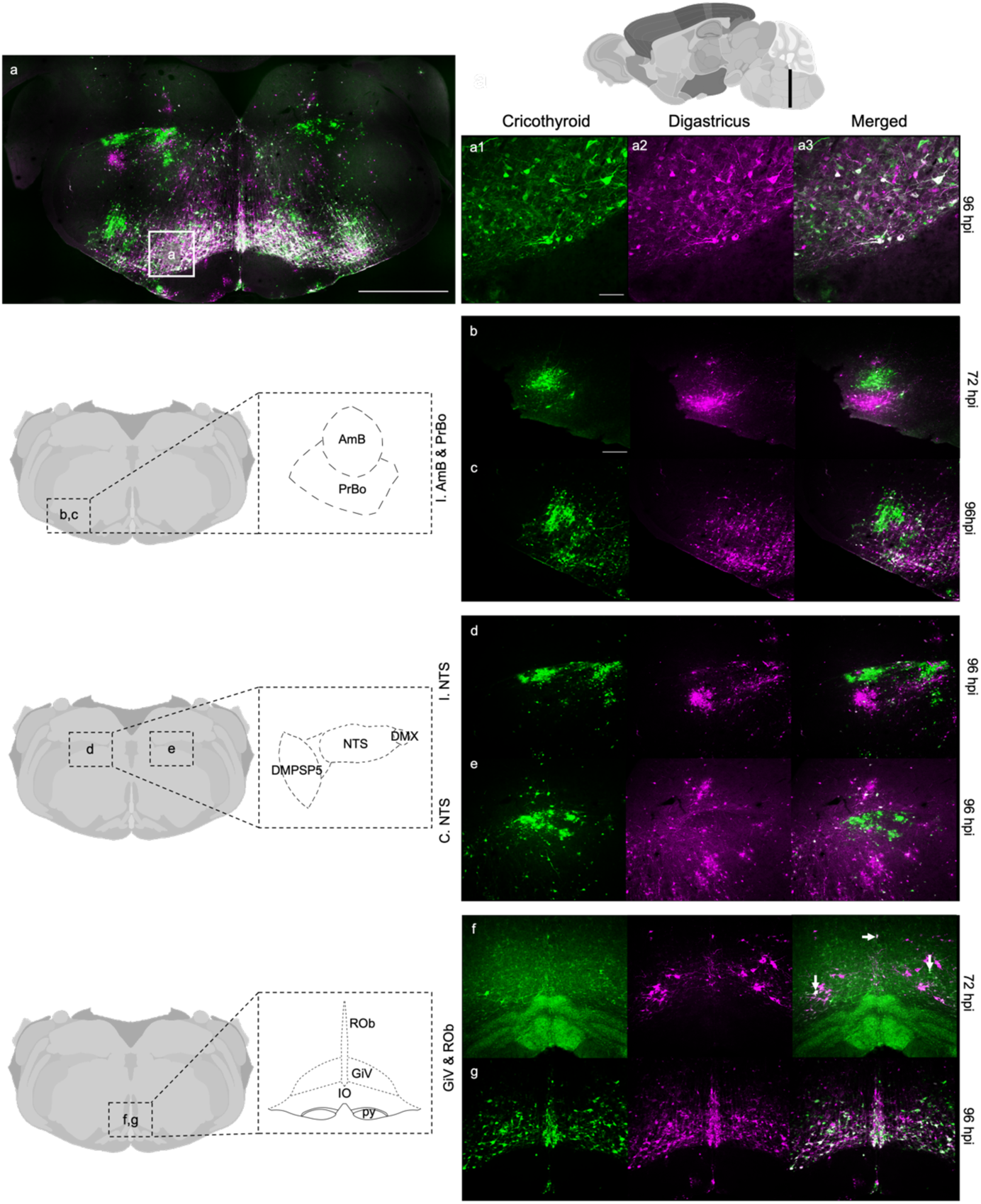
Hindbrain I. 2a, tile scan of the medulla. 2a1, higher magnification of colabeling, scale bar=100um. 2b, nucleus ambiguus at 72 hpi, scale bar=200um. 2c, nucleus ambiguus at 96hpi. 3d, ipsilateral nucleus of the solitary tract at 96hpi. 3e, contralateral nucleus of the solitary tract at 96 hpi. 2f, gigantocellular reticular nucleus and raphe obscurus at 72 hpi. 2g, gigantocellular reticular nucleus and raphe obscurus at 96hpi.

**Table 4:**
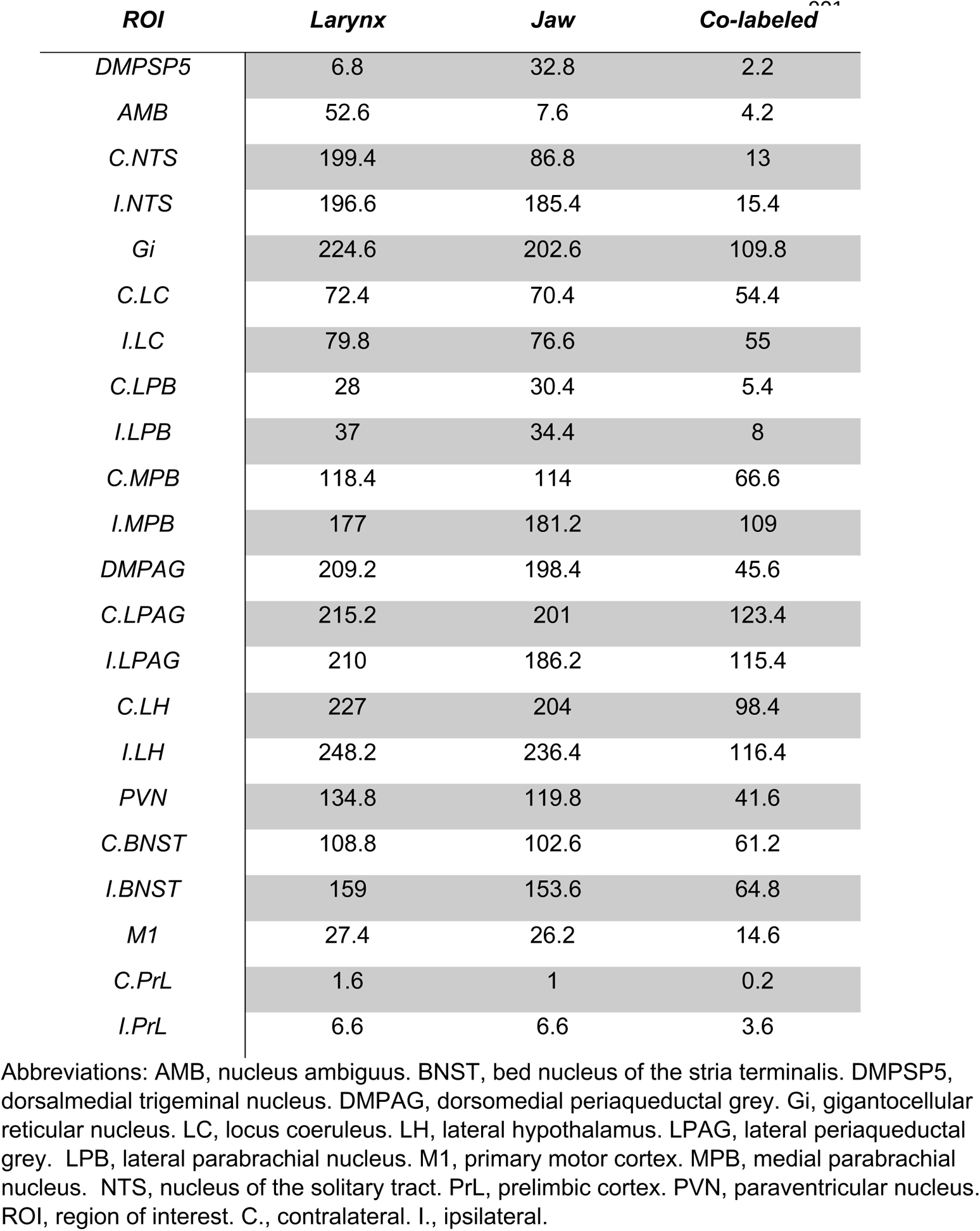
Average number of PRV-labeled neurons projecting to each structure and co-labeled.

| *ROI* | *Larynx* | *Jaw* | *Co-labeled*<sup>***</sup> |
| --- | --- | --- | --- |
| *DMPSP5* | 6.8 | 32.8 | 2.2 |
| *AMB* | 52.6 | 7.6 | 4.2 |
| *C.NTS* | 199.4 | 86.8 | 13 |
| *I.NTS* | 196.6 | 185.4 | 15.4 |
| *Gi* | 224.6 | 202.6 | 109.8 |
| *C.LC* | 72.4 | 70.4 | 54.4 |
| *I.LC* | 79.8 | 76.6 | 55 |
| *C.LPB* | 28 | 30.4 | 5.4 |
| *I.LPB* | 37 | 34.4 | 8 |
| *C.MPB* | 118.4 | 114 | 66.6 |
| *I.MPB* | 177 | 181.2 | 109 |
| *DMPAG* | 209.2 | 198.4 | 45.6 |
| *C.LPAG* | 215.2 | 201 | 123.4 |
| *I.LPAG* | 210 | 186.2 | 115.4 |
| *C.LH* | 227 | 204 | 98.4 |
| *I.LH* | 248.2 | 236.4 | 116.4 |
| *PVN* | 134.8 | 119.8 | 41.6 |
| *C.BNST* | 108.8 | 102.6 | 61.2 |
| *I.BNST* | 159 | 153.6 | 64.8 |
| *M1* | 27.4 | 26.2 | 14.6 |
| *C.PrL* | 1.6 | 1 | 0.2 |
| *I.PrL* | 6.6 | 6.6 | 3.6 |

Abbreviations: AMB, nucleus ambiguus. BNST, bed nucleus of the stria terminalis. DMPSP5, dorsalmedial trigeminal nucleus. DMPAG, dorsomedial periaqueductal grey. Gi, gigantocellular reticular nucleus. LC, locus coeruleus. LH, lateral hypothalamus. LPAG, lateral periaqueductal grey. LPB, lateral parabrachial nucleus. M1, primary motor cortex. MPB, medial parabrachial nucleus. NTS, nucleus of the solitary tract. PrL, prelimbic cortex. PVN, paraventricular nucleus. ROI, region of interest. C., contralateral. I., ipsilateral.

The nucleus of the solitary tract (NTS) became infected by both viruses at 84h, with few double-labelled neurons (Figure 2a, 2d, Table 4). Cricothyroid staining is bilateral in the NTS, while the digastricus stain is ipsilateral (Figure 2d,e). The infection of the NTS is extensive, spanning from the recess of the locus coeruleus to bregma −7.50mm (Franklin and Paxinos 2007), the caudal limit of our sample. Adjacent to the NTS, the ipsilateral dorsomedial spinal trigeminal nucleus (DMPSP5) labels for digastricus but not cricothyroid virus (Figure 2d). The DMPSP5 infection is restricted to the rostral ∼120μm of the nucleus, reflecting previous reports of its functional heterogeneity (Iv, Cheng, Takatoh, Han, & Wang, 2014).

The gigantocellular reticular nucleus (Gi) is a large, doubly-labelled region (Figure 2a, 2f, 2g). Together with the lateral paragigantocellular reticular nucleus (LPGi) and the raphe obscurus (ROb), these contiguous structures begin to be double-labelled at 72 hpi (Figure 2f). Unlike the nucleus ambiguus or DMPSP5, these nuclei are stained bilaterally from their first detectable infections (Figure 2g).

At the equivalent of coronal sections corresponding to bregma −5.70mm in mice (Franklin and Paxinos 2007), we observe extensive co-labelling in the locus ceruleus (LC) beginning at 84 hours (Figure 3a, 3b, 3c). To confirm the identity of the LC, we performed tyrosine hydroxylase (TH) immunolabeling on an alternate series of sections and found extensive triple labeling of TH, cricothyroid, and digastricus PRV infections (Figure 14a).

**Figure 3.**
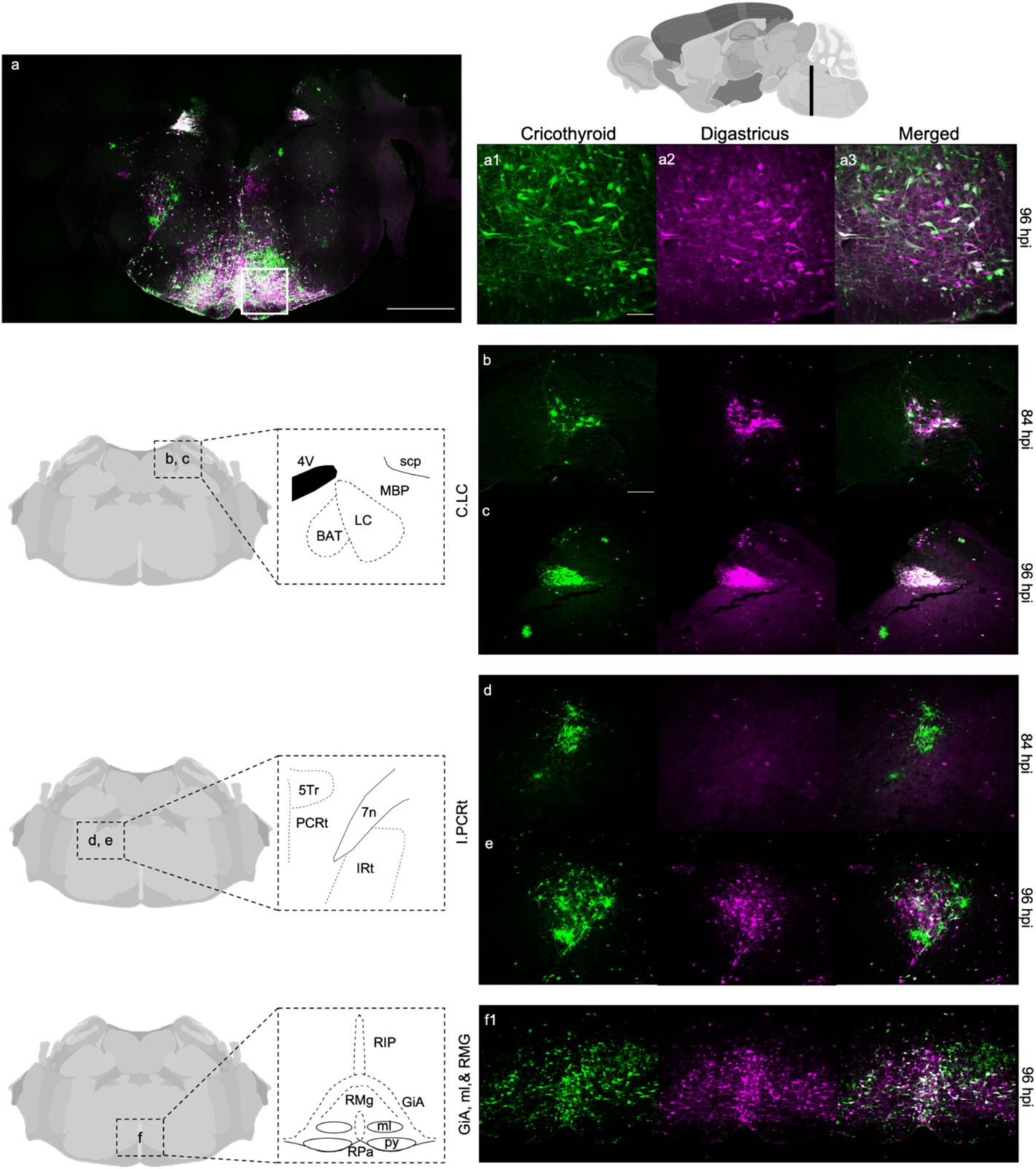
Hindbrain 2. 3a, tile scan of the pons, scale bar=1mm. 3a1, high magnification of co-labeling, scale bar=100um. 3b, locus coeruleus at 84 hpi, scale bar = 200μm 3c, locus coeruleus at 96 hpi. 3d, ipsilateral parvicellular reticular nucleus at 84 hpi. 3e, ipsilateral parvicellular reticular nucleus at 96 hpi. 3f, gigantocellular reticular nucleus, alpha part and raphe magnus nucleus at 96 hpi.

**Figure 14.**
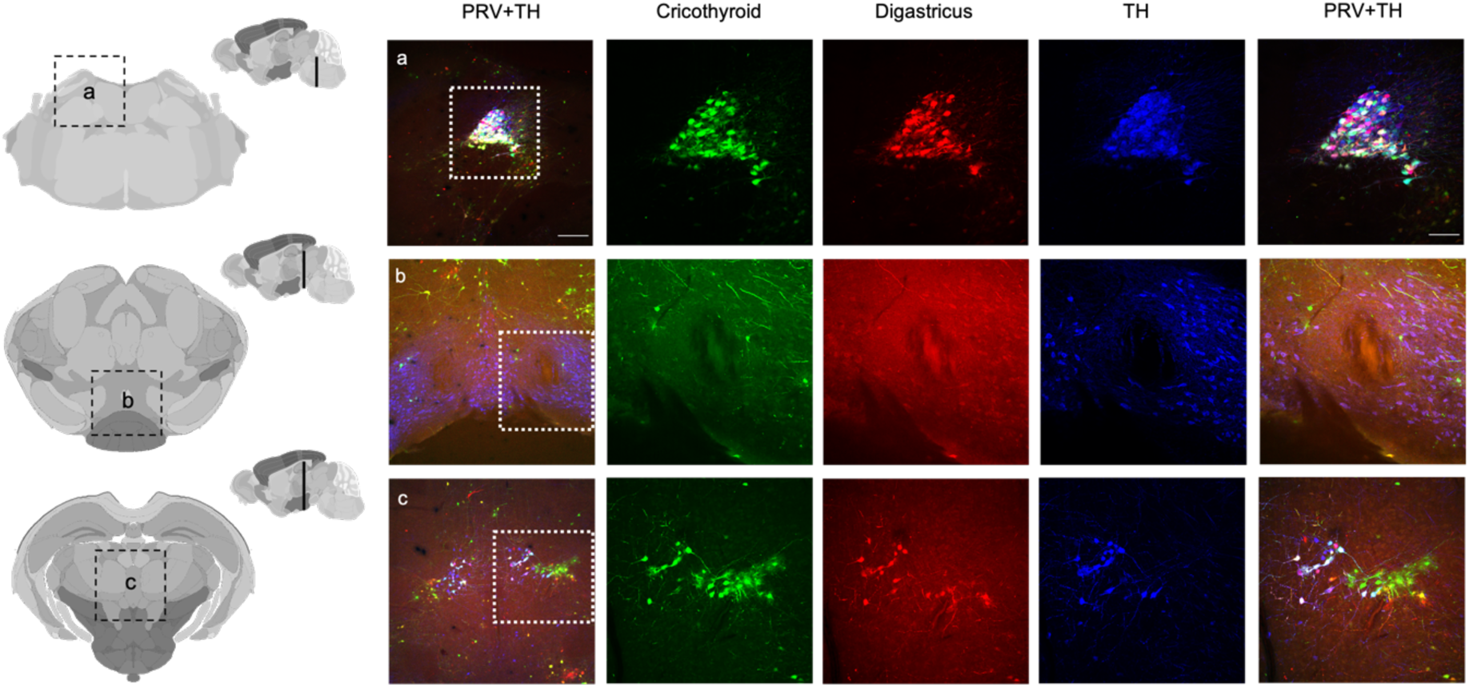
. Tyrosine hydroxylase triple labeled structures.

The parvicellular reticular nucleus (PCRT) is double-labelled, with the cricothyroid label emerging at 84h and the digastricus at 96h (Figure 3a-e). This infection may also include the adjacent intermediate reticular nucleus (IRT), but the boundaries between PCRT and IRT are not readily discernible.

As mentioned above, the gigantocellular reticular formation (Gi) extends rostrally into the plane that contains PCRT (Figure 3a, 3f), coincident with the beginning of the pyramidal tract and just rostral to the inferior olive, a level we interpret as Gi alpha (GiA). At this level, the Gi is contiguous with the caudal regions of the raphe magnus nucleus (RMg) at the equivalent of Franklin and Paxinos (2007) bregma −5.30 (Figure 3f).

At the level of the RMg, we note the primary motorneuron pool for the anterior digastricus, a subdivision of the motor trigeminal nucleus (5N) known as 5ADi. Beginning at 72h, it is infected by the digastricus PRV alone. The double-labelled neurons of RMg extend past the rostral boundary of the Gi (Figure 4a). The label is ipsilateral and remains specific to the digastricus-targeted PRV through 96h (Figure 4b, 4c). Outside of 5ADi, 5N is sparsely but specifically infected by the virus targeted to the digastricus (Figure 4c).

**Figure 4.**
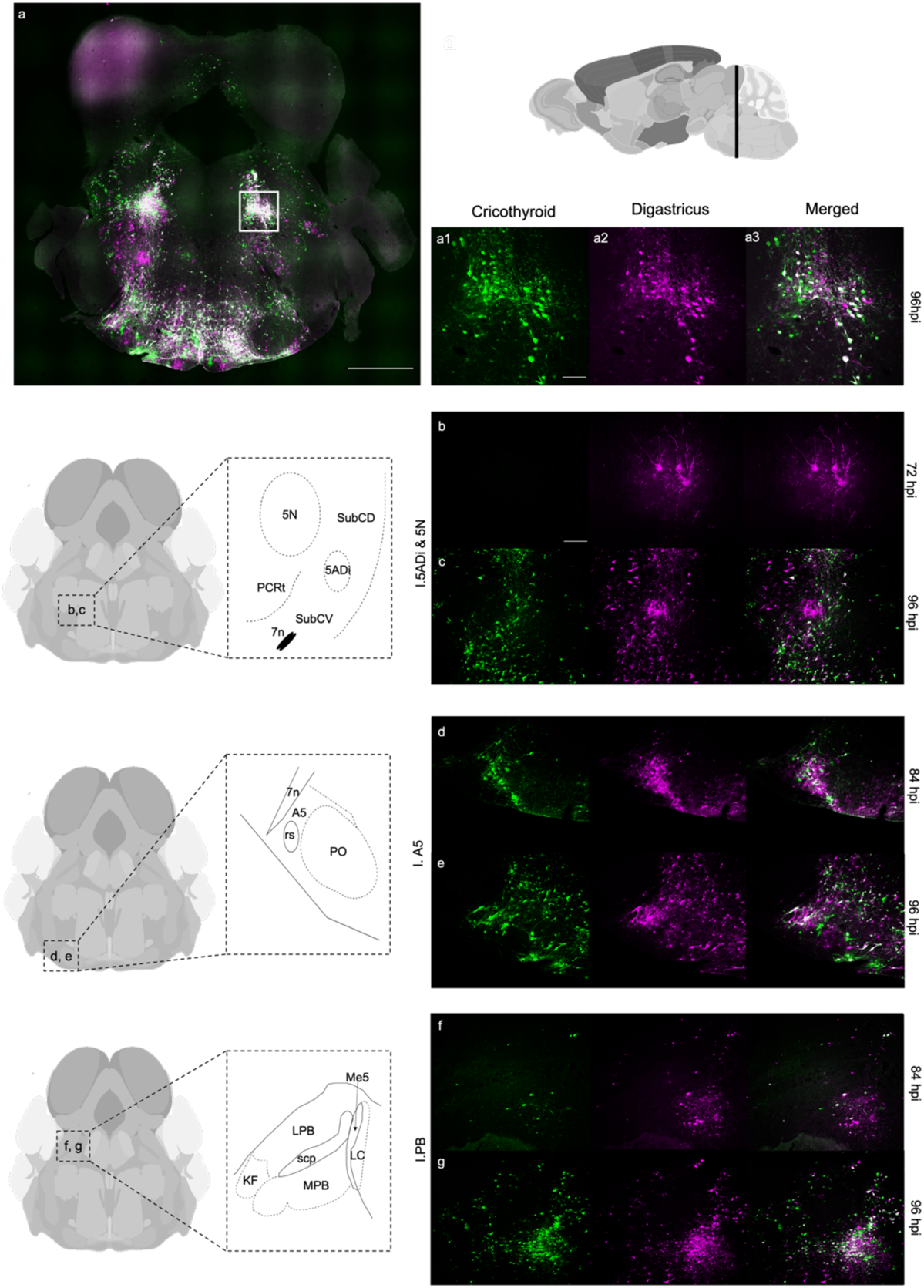
Hindbrain 3. 4a, tile scan of caudal portion of the midbrain, scale bar=1mm. 4a1, higher magnification of co-labeling, scale bar=100um. 4b, motor trigeminal nucleus, anterior digastric portion at 72 hours scale bar=200um. 4c, motor trigeminal nucleus, anterior digastric portion at 96 hours. 4d, Ipsilateral A5 cell population at 84 hours. 4e, Ipsilateral A5 cell population at 96 hours. 4f, ipsilateral parabrachial nucleus at 84 hours. 4g, ipsilateral parabrachial nucleus at 96 hours.

Just lateral to the paraolivary nucleus, we find another catecholaminergic site, A5 cells, are double-labelled at both the 84 and 96 hpi (Figure 3d, 3e). At 84 hours, the structure is more clearly delineated by co-labeling than at 96 hours (Figure 3d, 3e).

Lateral and ventral to the periaqueductal grey (PAG) and the aqueduct, we observe heterogeneous labelling among divisions of the parabrachial nucleus (PBN). At 84 hours, double-labelled cells arise in the medial parabrachial nucleus (MBP) while the lateral portion does not show substantial infection (Figure 4f). At 96 hours the MPB is densely co-labeled (Figure 4g), the lateral PBN has both labels, but few neurons are doubly labeled.

Regions of brainstem evident at the level of the PAG, specifically the PBN and the RMg, show similar patterns of double-labelling and timing as observed more caudally (Figures 4, 5), with sparse labeling at 84h and strong labeling at 96h. The pontine reticular nucleus (PnO) exhibits a similar pattern of infection, with sparse double-label evident at 84h, and more extensive staining at 96hpi (Figure 5f, 5g).

**Figure 5.**
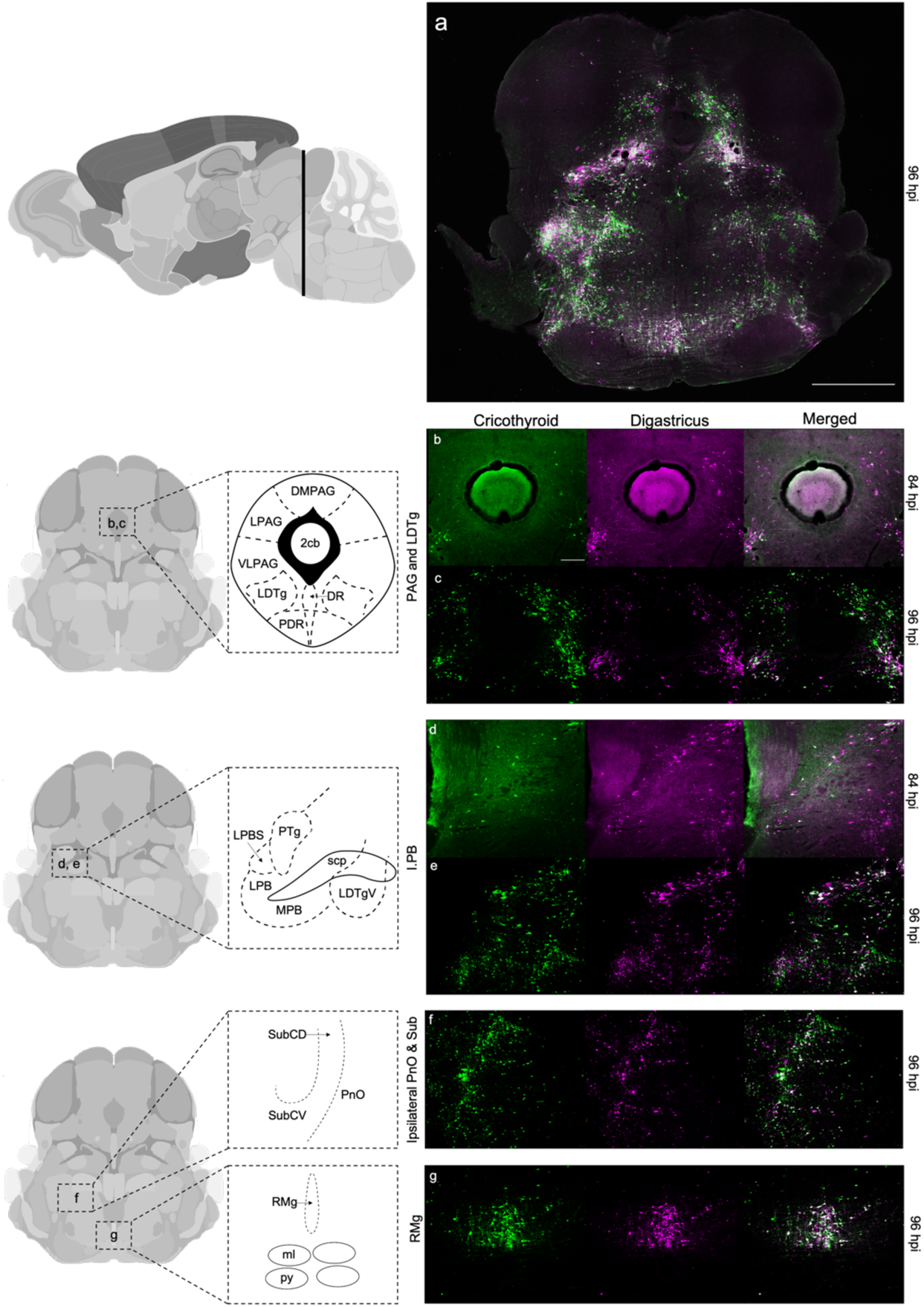
Midbrain 1. 5a, tile scan of the midbrain, scale bar=1mm. 5b, periaqueductal grey and lateral tegmentum at 84 hpi, scale bar=200um. 5c, periaqueductal grey and lateral tegmentum at 96 hpi. 5d, ipsilateral parabrachial nucleus at 84hpi. 5e, ipsilateral parabrachial nucleus at 96hpi. 5f, pontine reticular nucleus and subceuruleus at 96hpi. 5g, raphe magnus at 96hpi.

### Midbrain

The PAG, a canonical structure in the regulation of vocalization, shows substantial heterogeneity of labelling (Figures 5–9). Unlike regions of the brainstem, there is no labeling evident at 72h. At the caudal boundary of the PAG, where the cerebellar vermis appears in the aqueduct (Figure 5a), we see double-labelled cells appear in the lateral PAG (LPAG) and ventral lateral PAG (VLPAG) beginning at 84hpi (Figure 5b, 5c). There seems to be a lack of labeling of the dorsal lateral PAG (dlPAG) altogether, and a late arrival of labelling (96hpi) in the dorsal medial PAG (dmPAG). At 96hpi, labeling and co-labeling of the viruses is contiguous between the VLPAG and the lateral dorsal tegmentum LDtg (Figure 5c). The same pattern of labelling is evident rostrally (Figures 6–8).

**Figure 6.**
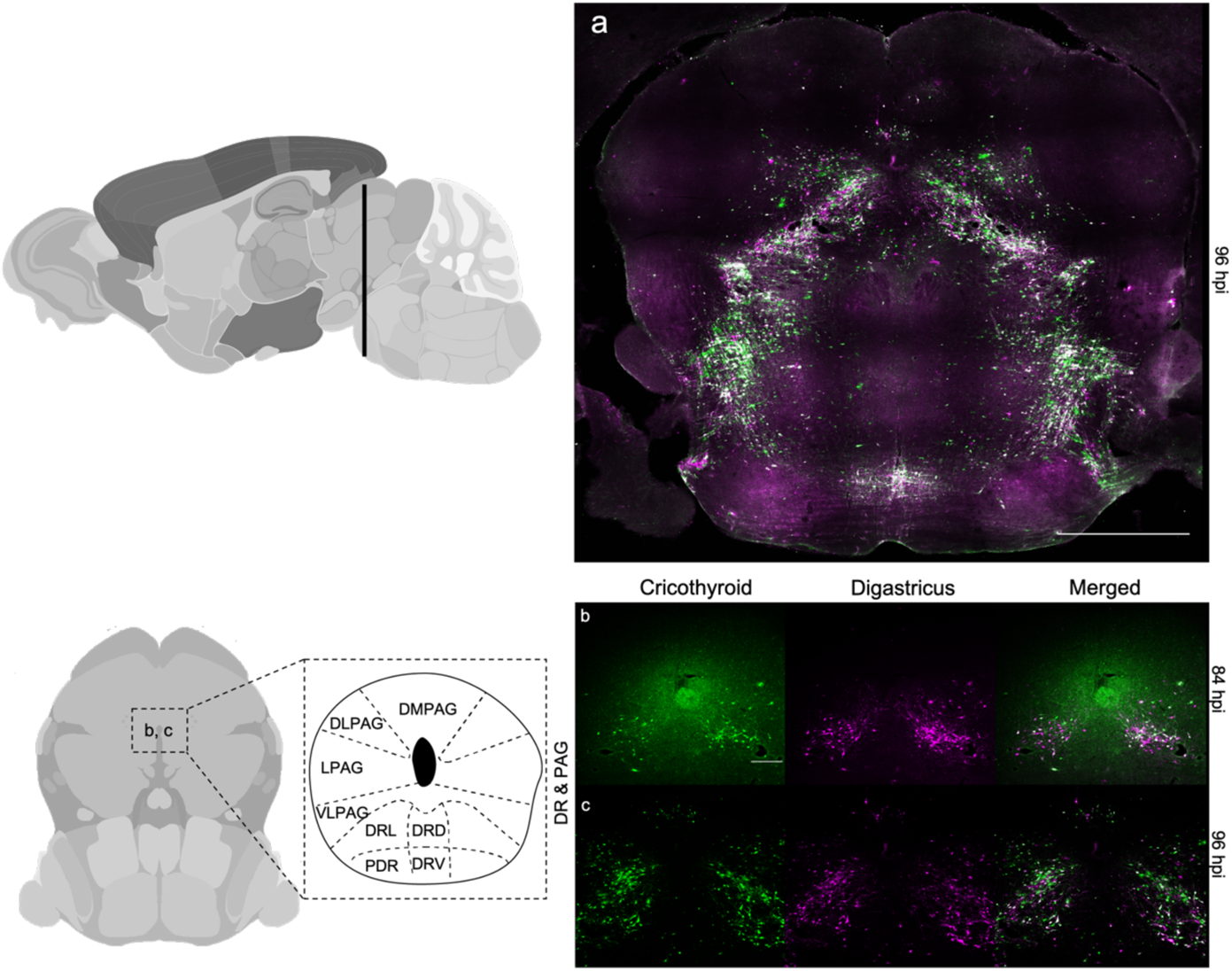
Midbrain 2. 6a, tile scan of the midbrain, scale bar=1mm. 6b, periaqueductal grey and dorsal raphe at 84 hpi, scale bar=200um. 6c, periaqueductal grey and dorsal raphe at 96 hpi.

**Figure 7.**
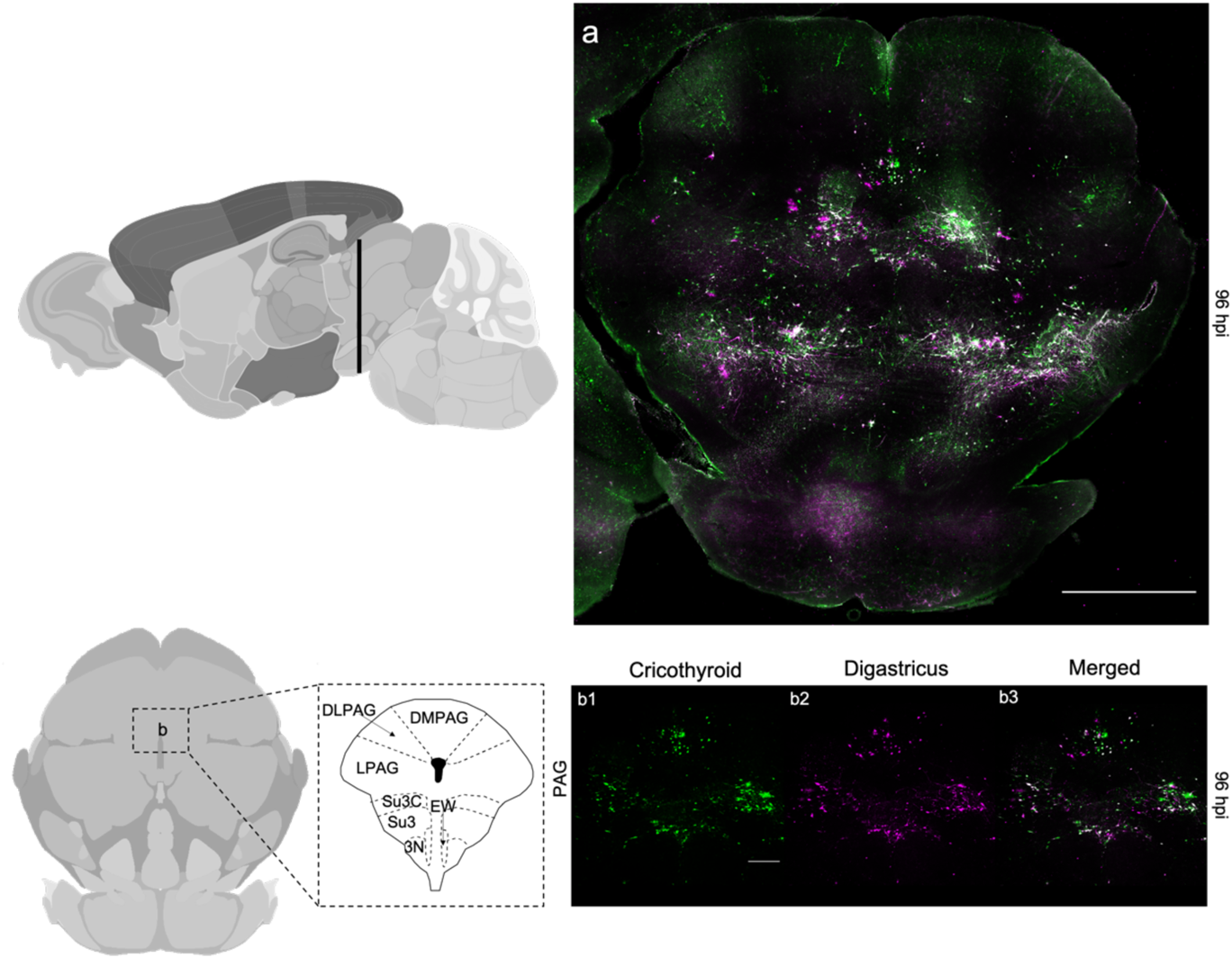
Midbrain 2. 7a, tile scan of the midbrain, scale bar=1mm. 7b, periaqueductal grey at 96hpi, scale bar=200um.

**Figure 8.**
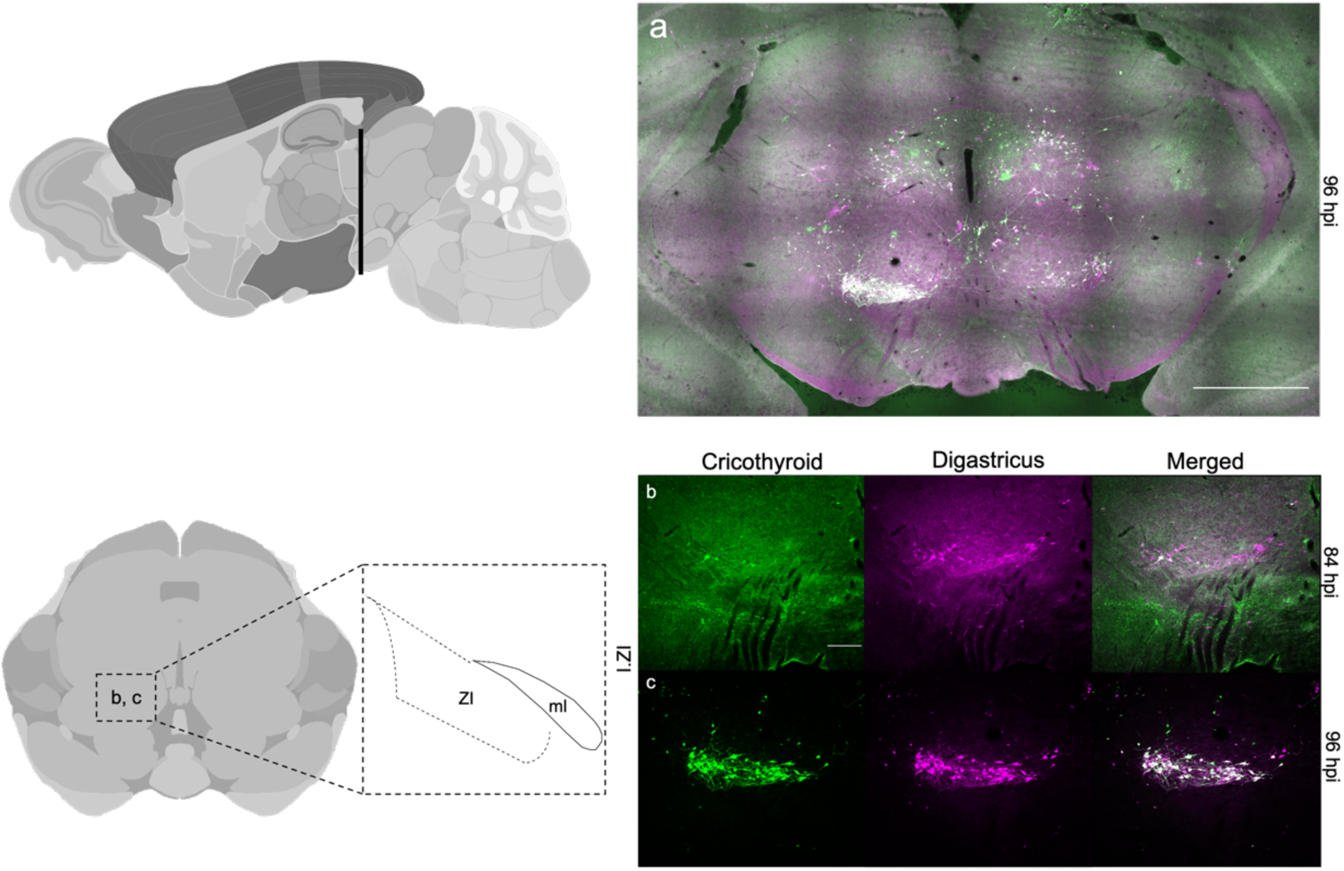
Thalamus 1. 8a, tile scan of the thalamus, scale bar=1mm. 8b, zona incerta at 84hpi. 7c, scale bar=200um, zona incerta at 96hpi.

**Figure 9.**
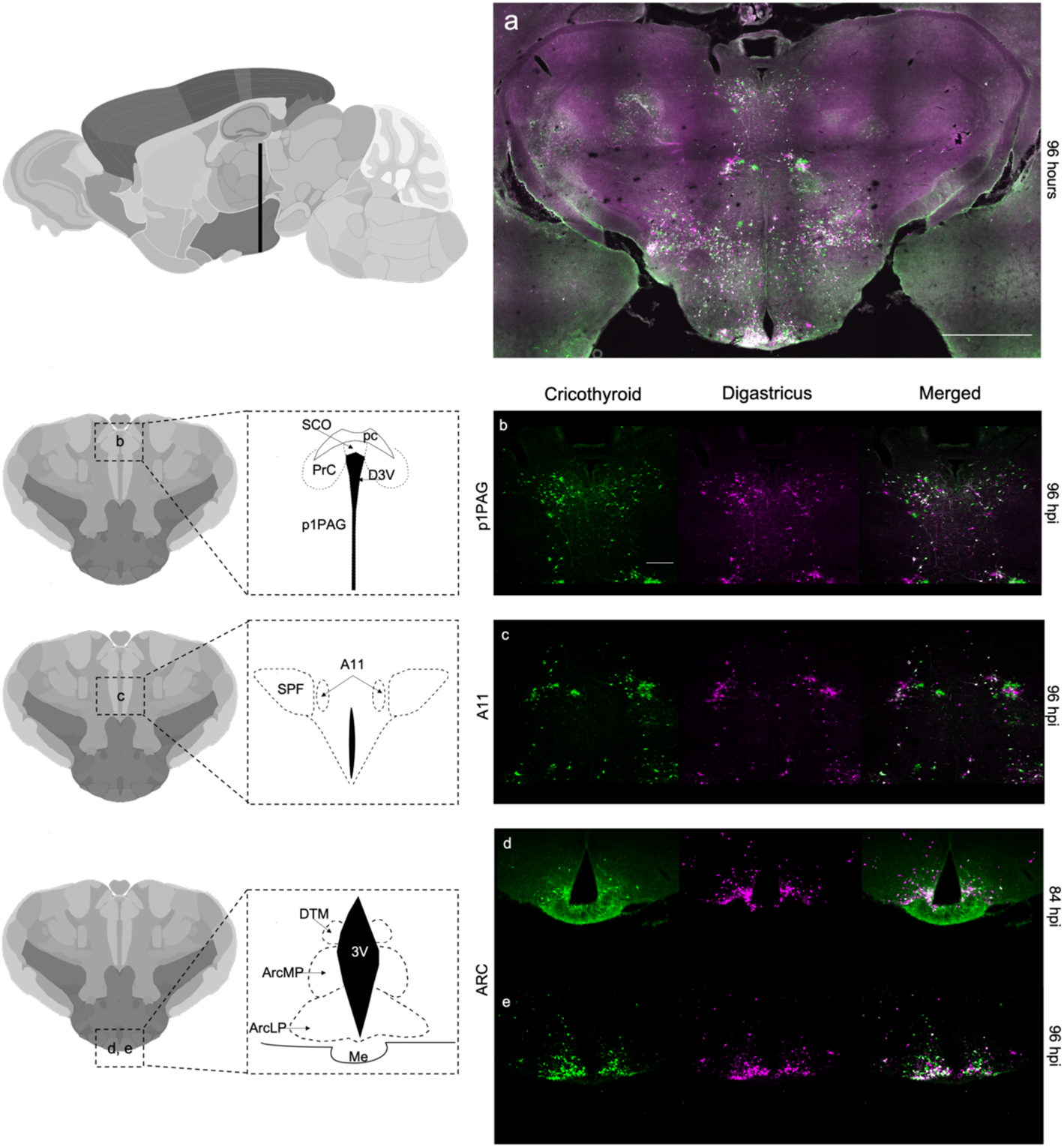
Thalamus 2. 9a, tile scan of the thalamus, scale bar=1mm. 9b, p1 periaqueductal grey at 96hpi, scale bar=200um. 9c, A11 cell population at 96hpi. 9d, arcuate nucleus at 84hpi. 93, arcuate nucleus at 96hpi.

At the equivalent of bregma −3.00mm (Paxinos and Franklin 2007), the zona incerta (ZI) exhibits digastricus label at 84hpi, and is dually labelled at 96 hours (Figure 8b, 8c).

At the rostral end of the PAG, the p1PAG is co-labeled beginning at the 96-hour time point (Figure 9a, 9b). At this same level, the A11 dopaminergic cell group is also co-labeled for cricothyroid and digastricus PRV beginning at 96hpi (Figure 9c). The catecholaminergic identity of the putative A11 group was confirmed by TH labeling (Figure 14c); the group of TH+ and PRV-double-labelled cells seemed more expansive than reported in Franklin and Paxinos (2007).

### Hypothalamus and amygdala

The arcuate nucleus (ARC) is co-labeled by the two viruses at both 84 and 96 hours (Figures 9, 10). The co-labeling was specific mostly to the lateral posterior portion with no labeling in the median eminence. At the caudal boundaries of the arcuate nucleus labeling, we also observe co-infection of the dorsomedial hypothalamus (DM) (Figure 10a, 10b) and the lateral hypothalamus (LH) (Figure 10c, d) at both 84 and 96 hours. We note only a small portion of the DM was labelled, and none of the ventral medial hypothalamus (VMH).

**Figure 10.**
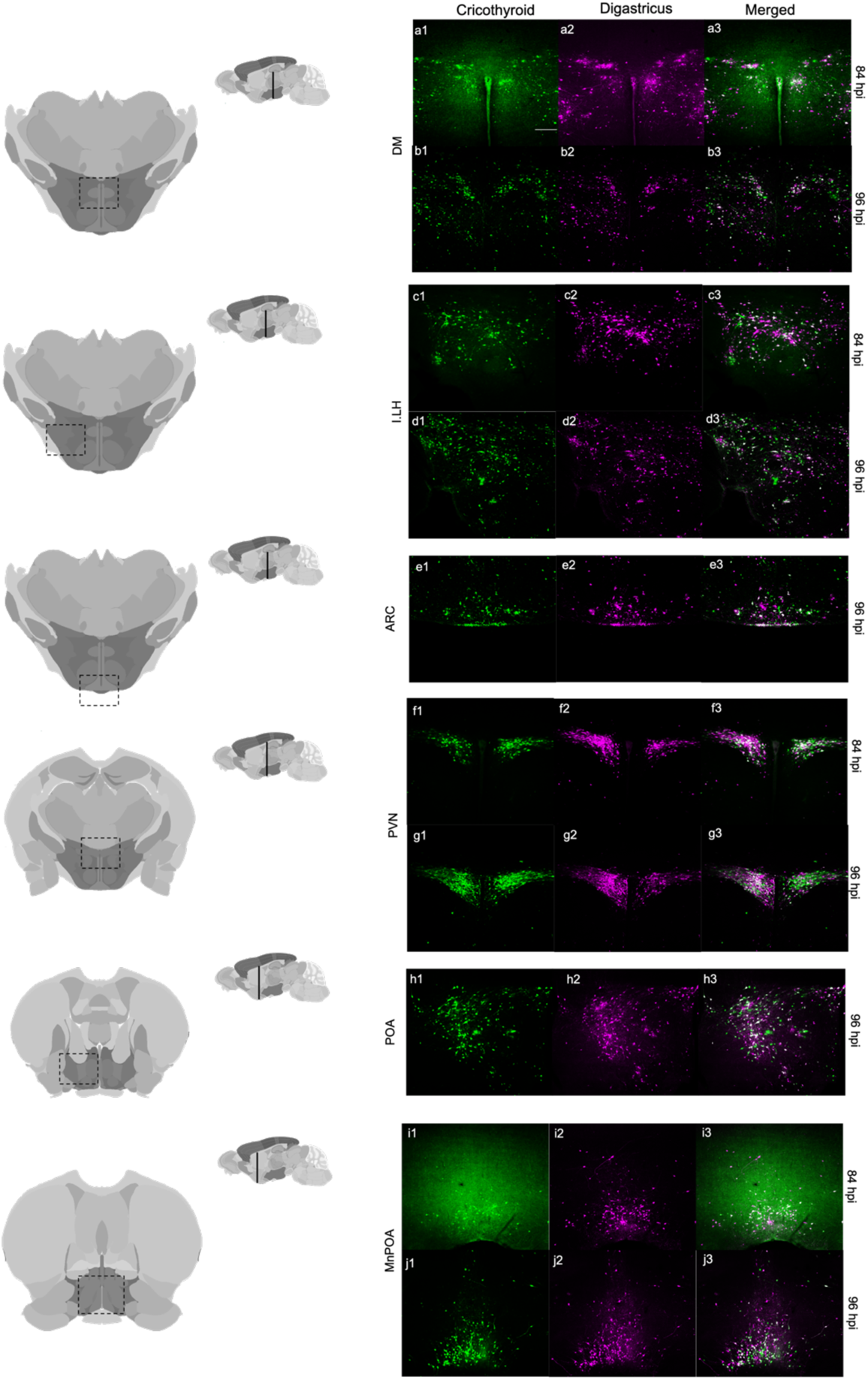
Hypothalamus. 10a, dorsomedial hypothalamus at 84 hpi, scale bar=200um. 10b, dorsomedial hypothalamus at 96hpi. 10c, ipsilateral lateral hypothalamus at 84hpi. 10d, ipsilateral lateral hypothalamus at 96hpi. 10e, arcuate nucleus at 96hpi. 10f, paraventricular nucleus at 84hpi. 10g, paraventricular nucleus at 96 hpi.10h, ipsilateral medial preoptic area at 96hpi. 10i, median preoptic area at 84hpi. 10j, median preoptic area at 96hpi.

Like the ARC, LH, and parts of the DM, the paraventricular nucleus of the hypothalamus (PVN) exhibits substantial double-labeling beginning at 84hpi (Figure 10f, 10g). Rostral to the PVN, the medial preoptic area (MPOA), as well as the median preoptic area (MnPOA), are strongly co-labeled beginning at 84hpi (Figure 10h,i), and more densely at 96hpi (Figure 10j).

The central amygdala (CeA) exhibits double-labelling in its medial portion beginning at 84hpi (Figure 11a) and expanding laterally into the remainder of the CeA at 96hpi (Figure 11b). Similarly, medial and rostral to the CeA, the extended amygdala (EA) begins to be co-labeled at 84 hours (Figure 11c) although the initial label is from the digastricus inoculation. At 96 hours, the EA is sparsely co-labeled (Figure 11d). Lastly, in the bed nucleus of the stria terminalis (BNST), co-labeling emerges at 96hpi (Figure 11e). At 96hpi, we observe a few double-labelled neurons in the lateral septum (not shown).

**Figure 11.**
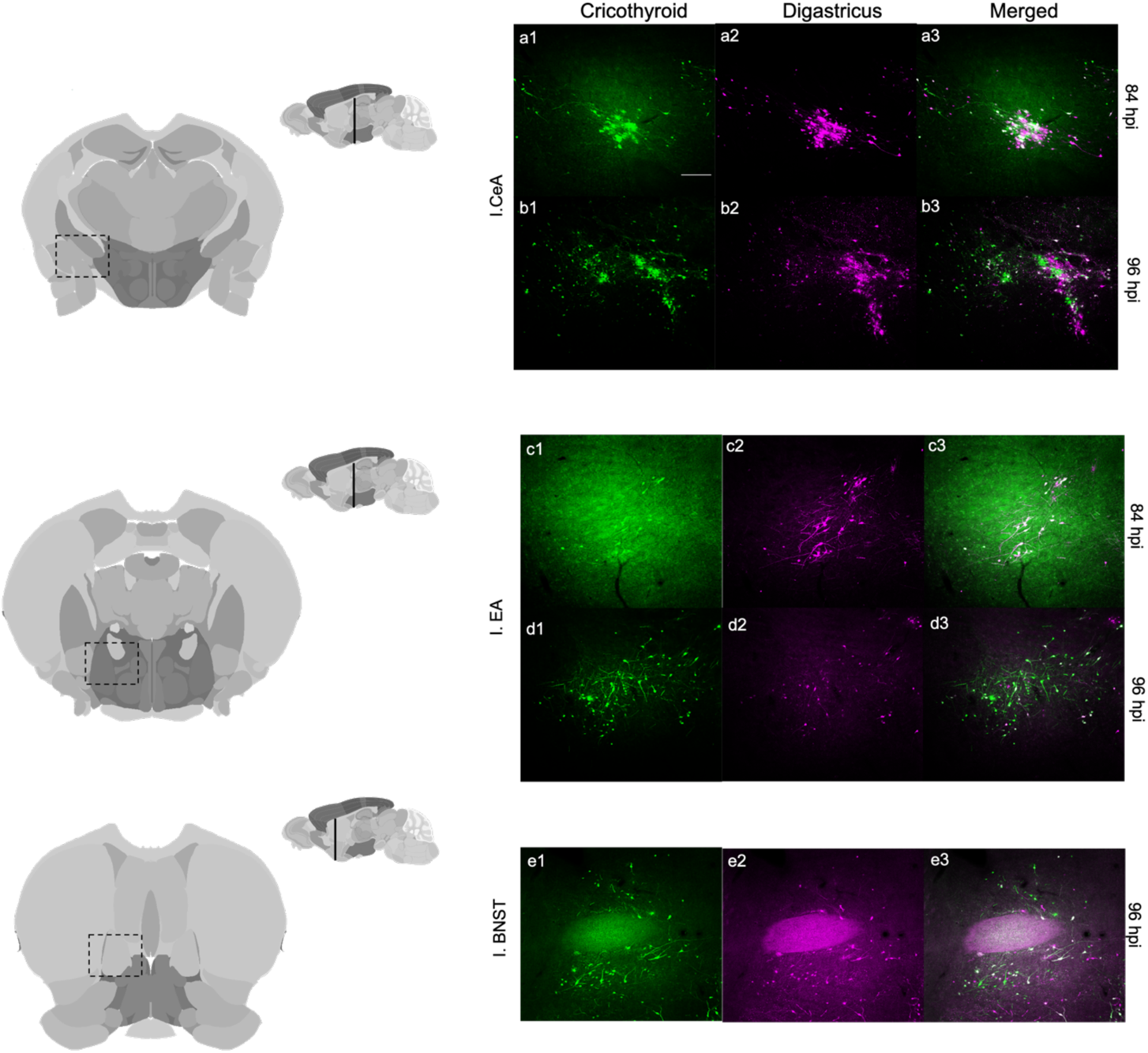
Amygdala. 11a, central amygdala at 84hpi, scale bar=200um. 11b, central amygdala at 96hpi. 11c extended amygdala at 84hpi. 11d, extended amygdala at 96hpi. 11e, ipsilateral bed nucleus of the stria terminalis at 96hpi.

### Cortex

We observe PRV infection in three distinct cortical regions in the *S.teguina* brain. At the caudal portions of the midbrain where the aqueduct appears, the ventral intermediate portion of the entorhinal cortex is labeled by virus targeted to the cricothyroid (Figure 12a). This bilateral pattern of infection appears at 96 hours. Near the bregma, we observe co-labeling in M1 motor cortex, corresponding to a region previously described as the mouse “laryngeal motor cortex” (Arriaga, 2012). This infection is strictly contralateral and emerges at 96hpi (Figure 12b). Lastly, we find a region in the caudal prelimbic cortex (PrL) that is co-labeled by the two viruses. At 84 hours this area is sparsely infected by the digastricus virus (Figure 12c) and at 96 hours, the overall infection is still sparse, but neurons are doubly labelled with both viruses (Figure 12d).

**Figure 12.**
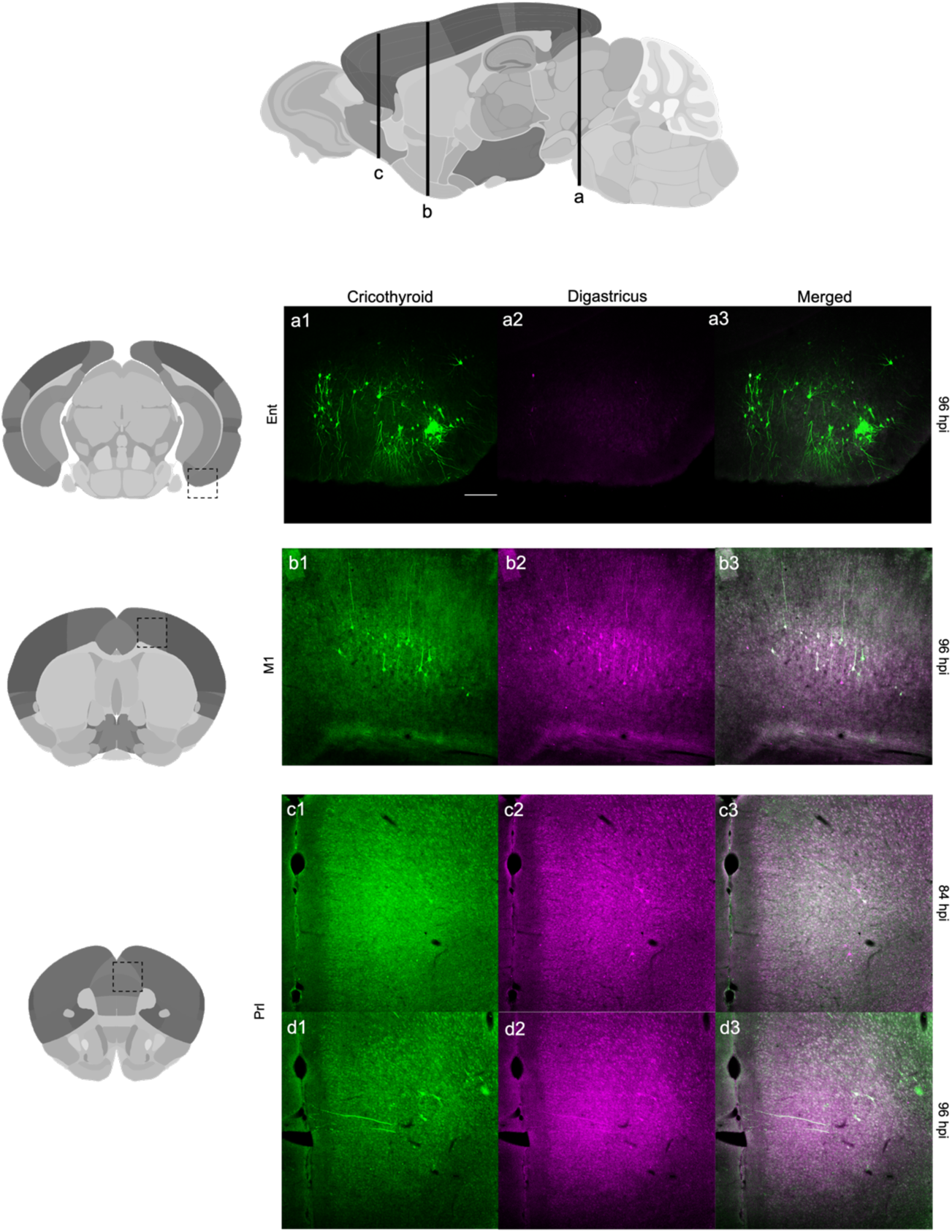
Cortex. 12a, contralateral entorhinal cortex at 96hpi, scale bar=200um. 12b, primary motor cortex at 96hpi. 12c, prelimbic cortex at 84hpi and 96hpi.

### Quantification of singly and doubly labeled neurons

To describe the extent of labelling for each virus, as well as the relative fraction of neurons that are co-labelled, we chose a subset of regions that spanned from spinal cord to limbic cortex (Table 3). We report both the average number of cells counted (Table 4) and the percentage co-labeled cells projecting to one structure or the other (Figure 13). One structure we report counts from in Table 4 is excluded from our description of percentage of co-labelled cells (PrL) due to the low number of infected neurons evident at 96hpi (Table 4).

**Figure 13.**
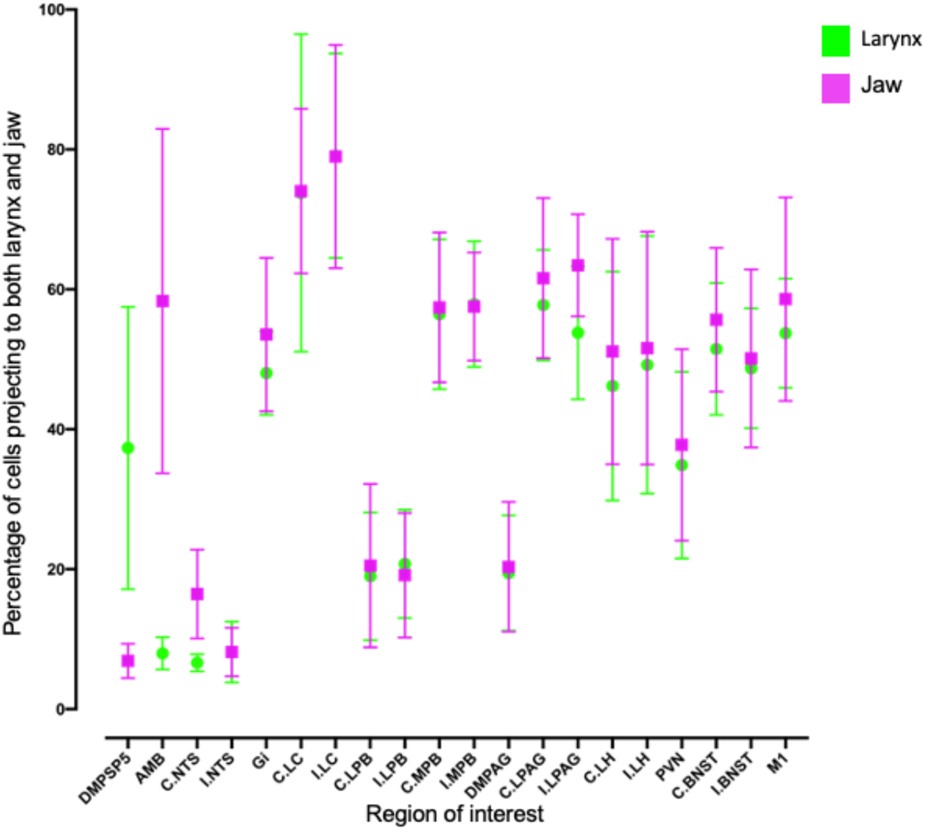
. Quantification and variation in co-labeled cells projecting to cricothyroid (larynx) and anterior digastricus (jaw) across regions of interest analyzed. Error bars=standard deviation. Abbreviations: AMB, nucleus ambiguus. BNST, bed nucleus of the stria terminalis. DMPSP5, dorsalmedial trigeminal nucleus. DMPAG, dorsomedial periaqueductal grey. Gi, gigantocellular reticular nucleus. LC, locus coeruleus. LH, lateral hypothalamus. LPAG, lateral periaqueductal grey. LPB, lateral parabrachial nucleus. M1, primary motor cortex. MPB, medial parabrachial nucleus. NTS, nucleus of the solitary tract. PVN, paraventricular nucleus. ROI, region of interest. C., contralateral. I., ipsilateral.

We find for regions containing motor neurons (nucleus ambiguus) or muscle-specific sensory neurons (DMPSP5), staining is overwhelmingly ipsilateral and, even at 96hpi, few neurons are double-labelled. The NTS, LPB, and the dmPAG are bilaterally infected by PRV targeted to both muscles, but have very low rates of co-infection (<20%). In contrast, the Gi, LC, MPB, and LPAG had high levels of double-labelled neurons (>40%). All measured forebrain structures had comparably high levels of co-infection, including the uniquely contralateral infection of the M1 cortex, as well as the bilateral infection of BNST, LH, and PVN. Overall, the pattern of double-labelling seemed to be bimodally distributed with a subset of structures exhibiting strong but segregated staining, and others exhibiting high levels of double-labelling.

Among brain regions with bilateral expression, we found that rates of double-labelling were similar across structures, but the absolute numbers of neurons infected and co-infected were larger in the hemisphere ipsilateral to the targeted muscles (Table 4, Figure 13).

## Discussion

We used two isogenic pseudorabies viruses (PRV) to characterize the neural circuits that terminate in muscles of the jaw and larynx, essential effectors of vocalization in the singing mouse, *Scotinomys teguina*. This dual-virus approach allows us to identify circuits specific to each of these muscles, as well as identify individual neurons that are upstream of both muscles (Figure 1). From these data, we examine the extent of single- and double-labelled neurons throughout the brain. Lastly, we examine the time of arrival of these vectors (Figure 15). We compare these results to reports from other species to assess whether the pattern of connectivity is consistent with vocal circuits in other vertebrates. Although we report a variety of novel findings, overall our anatomical data are consistent with characterizations of forebrain structures implicated in vocalization in other mammals, and with limbic, midbrain, brainstem and spinal cord patterns described in vocal vertebrates more generally. We now discuss these findings in detail for related regions of interest.

**Figure 15.**
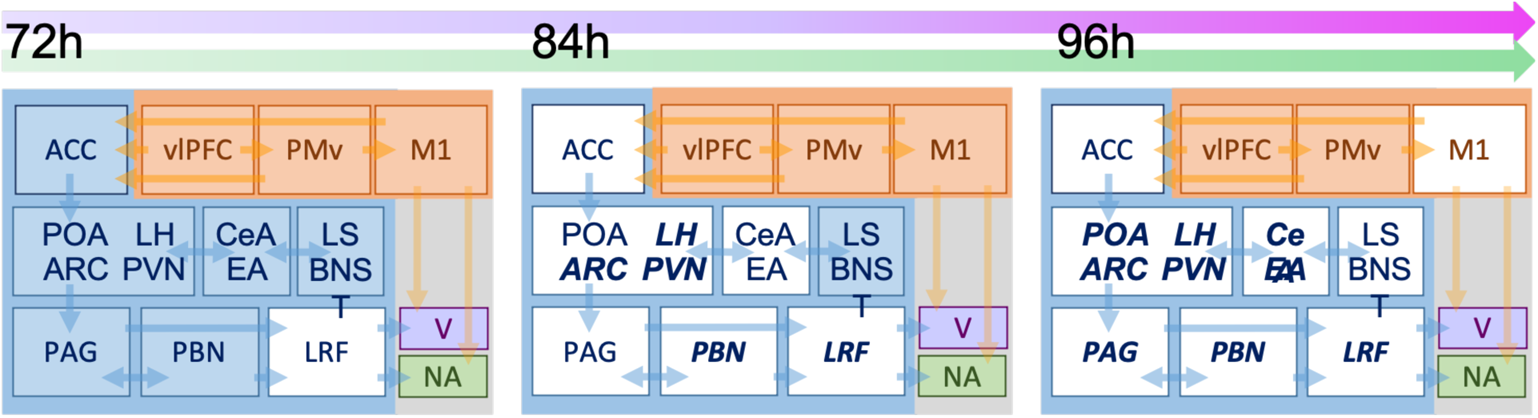
. Time course description of current PRV study. Blue represents the nodes of the primary vocal motor network as defined by Hage and Nieder 2016. Orange presents the nodes of the volitional articulatory motor network. Green and purple represent the infection by one of the strains of the virus. White represents evidence of co-labelling of both viruses. Bolded letters represent extensive infection of PRV in nuclei. Abbreviations: V, motor trigeminal nucleus and motor trigeminal nucleus, anterior digastric portion. ACC, anterior cingulate cortex as putatively defined by Bennet et al 2019 to be a sub compartment of the prelimbic cortex. ARC, arcuate nucleus. BNST, bed nucleus of the stria terminalis. CeA, central amygdala. EA, extended amygdala. LH, lateral hypothalamus. LS, lateral septum. M1, primary motor cortex. NA, nucleus ambiguus. PAG, periaqueductal grey. PBN, parabrachial nucleus. PMv, ventral premotor cortex. POA, preoptic area. RF, reticular formation. vlPFC, ventrolateral prefrontal cortex.

### Motor neurons and other singly labelled nuclei

The first detection of PRV in the CNS occurs as single-labelled cells in spinal cord compartments that correspond to motor neurons innervating either the larynx (cricothyroid) or the jaw (digastricus). The nucleus ambiguus (NA) is known to contain motor neurons innervating the intrinsic muscles of the larynx, including the cricothyroid. In our study, as in others, the NA is the first to be infected with virus injected into the cricothyroid and contains single-labelled neurons (Figure 2a, 2b; (Arriaga, 2012; Barrett, Bao, Miselis, & Altschuler, 1994.; Cassell, 2010; Hisa, 2016; Waldbaum, Hadziefendic, Erokwu, Zaidi, & Haxhiu, 2001)). Previous mapping studies suggest that the NA contains distinct fields of neurons specific to particular muscles in the throat. The cricothyroid compartment of the NA occupies a rostral portion of the nucleus, while the other intrinsic laryngeal muscles are more caudal (Hisa, Sato, Fukui, Ibata, & Mizuokoshi, 1984). Consistent with this observation, our cricothyroid infection appears rostrally, at the same rostral-caudal level as the inferior olive (Figure 2) (Barrett et al., 1994; Hisa, 2016).

Fewer studies have targeted the anterior digastric muscle, though for singing mice (and presumably many other species), it is an important vocal muscle (Okobi et al 2019). The motor neurons innervating the anterior digastricus form a distinct subnucleus (5ADi) that is a ventromedial compartment of the motor trigeminal nucleus (5N) (Franklin & Paxinos, 2007). In our data, we note an early, single-label ipsilateral infection of 5ADi that persists to 96 hours (Figure 4a, 4b). We also find that the ipsilateral motor nucleus of the trigeminal (5N) is sparsely infected Figure 4c). This is consistent with horseradish peroxidase (HRP) and PRV studies targeting the anterior digastric belly of the muscle of rats (Kang et al., 1999; Kemplay & Cavanagh, 1983).

In addition to motor neurons of the jaw and larynx, we also found a predominant ipsilateral single label of the spinal trigeminal nucleus oralis (DMPSP5) (Figure 2d, Table 4, Figure 13). This nucleus is considered a somatosensory nucleus, and also has been labelled in monosynaptic retrograde studies of the masseter (Iv et al., 2014). Given the importance of the anterior digastric muscle in the vocalizations of *S.teguina* (Okobi et al., 2019) specifically with gape and frequency modulation, we speculate this nucleus may provide sensory feedback on jaw movement during vocalization and other activities.

We find bilateral infection of the nucleus of the solitary tract (NTS) (Figure 2d, 2e), a pattern observed in previous work (Arriaga et al., 2015; Barrett et al., 1994; Hisa, 2016). The NTS is often considered upstream of motorneuron pools, as a primary premotor nucleus and as a second-order regulator of the nucleus ambiguus. We find that this structure and many of its sub-compartments are infected by PRV originating in the cricothyroid. A subset of neurons are labelled with PRV originating in the digastricus, but relatively few neurons are double-labelled (Table 4, Figure 13). This suggests the structure is heterogenous in function; although it seems positioned to influence both muscle groups, few cells seem equipped to coordinate both muscles. Its strong expression in the ipsilateral side by the digastric infection suggests independent modulation of jaw function. Functional tests of this region have implicated it in innate vocalization and tied it to the expiratory phase of breathing (Hernandez-Miranda et al., 2017).

### Putative central pattern generators

The *Scotinomys* song is a rhythmic and frequency-modulated trill that involves the coordination of larynx, jaw, and respiratory muscles to produce each note (Campbell et al., 2010; Okobi et al., 2019; Pasch et al., 2011b). The notes themselves are repeated and stereotyped, and acoustic changes over the course of the song are well-described by a polynomial function (Campbell et al., 2010). This highly rhythmic and repeated coordination of movements suggests the activity of one or more central pattern generators; we would expect such CPGs to be double-labelled bilaterally, to reside within the brainstem or spinal cord, and to be among the first structures doubly labelled.

Work on vocalization and orofacial pattern generation generally suggests that CPGs for vocal behavior would be located in the brainstem (Barlow, 2009; Bass, 2014; Bass & Remage-healey, 2008; Hage, 2010; Moore, Kleinfeld, & Wang, 2014; Rhodes, Yu, & Yamaguchi, 2007). Among the many CPGs implicated in orofacial movements (Moore et al 2016), Hage (2010) suggests five candidates for patterning of vocal behavior specifically. These include the parvicellular reticular nucleus (PCRt), pontine reticular nucleus (PnO), the nucleus retroambiguus (NRA), the nucleus raphe magnus (RMg), and the lateral paragigantocellular reticular (LPGi) formation.

Among these regions, the LPGi labelling is most consistent with our expectations of a CPG. It is labeled at the earliest timepoint (72hpi), largely double-labelled, and bilaterally infected (Figure 2a, 3a, 13, Table 4). The pattern of double-labelled neurons form a large medial structure that spans not only the LPGi, but extends throughout the gigantocellular reticular formation (Gi) and raphe obscurus (ROb) – structures that have been identified as orofacial pattern generators, but were not identified as putative vocal pattern generators (Moore et al. 2016, Hage 2010).

Among three other putative CPGs, we find labelling that seems inconsistent with expectations for a vocal pattern generator. For example, expression of the digastricus and cricothyroid labels in the PCRT is largely segregated by muscle (Figure 3d, 3e). Both the PnO and RMg exhibit double-labelling, but the viruses do not arrive in either structure until 84 hpi, a time point that coincides with expression in the hypothalamus and other upstream structures (Figure 5f, 5g). Lastly, the NRA is at the caudal boundary of our tissue sampling, and so we did not reliably obtain sections for examination; the literature supporting its role in vocalization (Tschida et al. 2019) suggests it would be worth examining this structure in a more targeted way.

From our current data, the LPGi and contiguous Gi and ROb would seem to be the strongest candidates for CPGs governing song production. However, there are a variety of models that describe how CPGs may control orofacial movements including vocalizations (Hage 2010, Stanek IV et al 2014), and it is possible that one or more of additional brainstem regions may influence patterning of the *Scotinomys* song.

### Neuromodulatory regions

We find several regions within the pons that are known to express either neuropeptides or catecholamines and exhibit robust double-labelling at 84hpi. We interpret these structures as neuromodulatory regions.

Our study points to a strong role of catecholamines in the modulation of vocal behavior in this species. The noradrenergic A5 and A6 (locus ceruleus, LC) populations are co-infected by the two PRV strains (Figure 3b, 3c, 4d, 4e). Indeed, LC exhibits an even higher degree of co-localization than the Gi (Figure 13, Table 4). The LC is the major center for norepinephrine synthesis in the brain (Samuels & Szabadi, 2008). It has major connections throughout the neuraxis (Szabadi, 2013) including the nucleus ambiguus, dorsal motor root of the vagus, and the gigantocellular reticular formation. The LC has been indirectly implicated in ultrasonic vocalizations (Hamed & Boguszewski, 2018) and is more broadly involved in wakefulness, arousal, and sensory processing (Aston-Jones & Cohen, 2005; Samuels & Szabadi, 2008). Interestingly, the LC influences perception of rodent vocalizations through its projections to the auditory cortex (Foote, Freedman, & Oliver, 1975; Martins & Froemke, 2015; Sara, 2009). Perhaps more closely related to the current study, the cell groups of A5 and A6 also play antagonistic roles in the regulation of respiratory rhythms (Guyenet, Koshiya, Huangfu, Verberne, & Riley, 1993; Hilaire, Viemari, Coulon, Simonneau, & Bévengut, 2004). The neural circuitry underlying breathing is obviously of broad relevance to vocalization in general (Barkan & Zornik, 2020), and in singing mice, each note of its song is accompanied by a short breath (Okobi, 2016; Pasch et al., 2011a)

We also observe co-infection in the dopaminergic A11 population of cells (Figure 9c). Like the noradrenergic populations, these neurons are known to have roles in auditory processing (Nevue, Felix 2nd, & Portfors, 2016) as well as motor functions (Koblinger et al., 2014). Interestingly, our dopaminergic results contrast with work that emphasizes the role of dopamine in other taxa (Saravanan, Hoffmann, Jacob, Berman, & Sober, 2019; Simonyan, Horwitz, & Jarvis, 2012), in that our TH/PRV triple-labeling revealed no PRV infection in either the ventral tegmental area or the substantia nigra pars compacta (Figure 14b).

The parabrachial nucleus (PBN) expresses a variety of neuropeptides, and is known for its role in vocal-respiratory interactions (Smotherman, Schwartz, & Metzner, 2010). The structure is topographically organized in terms of lateral, ventrolateral, and medial regions surrounding the superior cerebellar peduncle (Paxinos and Franklin 2007). In our study, the medial division has the highest abundance of double-labelled cells (Figure 13, Table 4). This division has reciprocal connections with the nucleus ambiguus (Herbert, Moga, & Saper, 1990; Núñez-Abades, Portillo, & Pásaro, 1990; Saper & Loewy, 1980) and receives inputs from much of the forebrain, including limbic regions that are canonical parts of the mammalian vocal circuit, such as hypothalamus, amygdala, anterior cingulate cortex (ACC) and laryngeal motor cortex (LMC)(Uwe Jürgens, 2002). Recordings in cats show that the medial PBN fires along with vocal and respiration behaviors (Farley, Barlow, & Netsell, 1992). In free-tailed bats (*Tadarida brasiliensis*), *c-fos* immunoreactivity is detected in the medial (and lateral) parabrachial nucleus after calling (Schwartz & Smotherman, 2011). It is hypothesized that the medial subdivision of the parabrachial nucleus modulates laryngeal activity for the purpose of vocalizations while the lateral is directed toward vocal-respiratory coupling (Smotherman et al., 2010). The extensive double-labelling apparent in our experiment suggest the parabrachial nucleus may more broadly coordinate vocal muscles (Figure 4g, 5e).

### The midbrain

The periaqueductal grey (PAG) is considered one of the most crucial centers for mammalian vocalization (Gruber-Dujardin, 2010; Uwe Jürgens, 2002). Multiple lines of evidence point to this large heterogeneous structure as one for “gating” of downstream vocal output (Esposito, Demeurisse, Alberti, & Fabbro, 1999; U Jürgens, 1994; Tschida et al., 2019). Anatomically, it stretches rostrally to the hypothalamus and caudally to the pontine nucleus of the brainstem (Gruber-Dujardin, 2010). The structure has been subdivided by Bandler and Keay (1996) along its longituditional axis into four columns: the dorsomedial PAG (dmPAG), dorsolateralPAG (dlPAG), lateral PAG (LPAG), and ventrolateral PAG (vlPAG) (Bandler & Keay, 1996; Kingsbury, Kelly, Schrock, & Goodson, 2011). These subdivisions are thought to serve different functional domains and promote different types of vocal output (Dujardin & Jürgens, 2006). Our data highlights distinctions among these subdivisions (Figures 5–9).

The vlPAG is consistently co-infected from the rostral to caudal axis of the PAG, and at 84hpi, it is the earliest infected of the PAG compartments. Among the remaining compartments, the LPAG has the higher co-infection (Figure 13). The dmPAG has robust infection by both viruses, but few doubly labelled cells (Table 4). Moreover, label in the dmPAG doesn’t arrive until 96hpi. Together, the data suggest the LPAG and vlPAG are more likely to regulate vocalization in *S.teguina*. This heterogeneity among PAG regions is consistent with tracing studies targeting the jaw and/or larynx of laboratory rodents (Bennett et al., 2019; Falkner et al., 2020; Fay & Norgren, 1997; Tschida et al., 2019). Moreover, among both mammals and birds, the LPAG is thought to be particularly important for vocalizations (Dujardin & Jürgens, 2006; Kingsbury et al., 2011).

### The hypothalamus

The hypothalamus is a major modulator of vocal behavior (Adkins-Regan, 2005; Hage, 2010; Hage & Nieder, 2016; Nieder & Mooney, 2020). In singing mice, hypothalamic areas such as POA, PVN, and LH all show medium to high levels of co-infection by the two strains of virus (Figure 13, Table 4). The POA and the PVN are intricately involved in courtship and aggressive behaviors (Wei et al., 2018), and are canonical regions of a social behavior network important across vertebrates (Connell & Hofmann, 2011; Goodson, 2005; Newman, 1999.). We find dense co-labeling in both the medial preoptic area and the median preoptic area (Figure 10h-j). The MPOA is implicated vocal and courtship behavior across vertebrates (Goodson & Bass, 2000; Schmidt, 1968), and in the generation of rodent ultrasonic vocalizations specifically (Gao, Wei, Wang, & Xu, 2019), Michael and Tschida et al 2019). Studies of squirrel monkeys implicate the dorsomedial hypothalamus (DM) in species-specific calls (Jurgens and Ploog 1970, Jurgens 1982), most likely due to its connections to the PAG (Dujardins and Jurgens 2005); we find doubly-labelled cells in the DM, but they are restricted to a subset of the nucleus (Figure 10a, b).

The ARC, PVN, and LH (Figure 9d, 9e, 10c-g) are interconnected nuclei that regulate energy balance in part through effects on feeding and other behaviors (Larsen et al 1994, Morton et al 2006, Stuber and Wise 2015). One of the many studies implicating these regions in energy balance is a double-PRV study showing viruses injected into liver and adipose tissue converge on neurons within these hypothalamic structures (Stanley et al 2010). The confluence of social and feeding circuits within the PVN in particular suggests that it is well-positioned to play a role in the modulation of vocal effort by energy balance, a pattern known in singing mice, as well as many other species (Burkhard et al 2018, Giglio and Phelps 2020, Morton 2017).

### The extended amygdala

The amygdala complex plays a central role in a diversity of motivated behaviors, many of which are social (Newman 1999). The lateral septum (LS), central amygdala (CeA) and bed nucleus of the stria terminalis (BNST) have all been implicated in vocalization across diverse taxa (see below). In *Scotinomys,* we find co-labelling from larynx and jaw in the CeA, BNST, and in the extended amygdala (*sensu Franklin and Paxinos 2007*). The EA and CeA both are co-infected at 84 hours (Figure 10a-d), while we did not detect double labeling in the BNST and LS until 96 hours (Figure 10e). Even at 96h, LS labeling was relatively sparse, suggesting it is upstream of amygdalar or hypothalamic structures. In lab mice, the EA and CeA inhibit the production of USVs through descending projections to the PAG (Michael and Tschida 2019). In contrast, electrical stimulation of the amygdala can elicit vocalizations in a variety of mammals, including the mustache bat (Ma and Kanwal 2014), guinea pig (Green et al 2018), domestic pig (Manteuffel et al 2007) and squirrel monkey (Jurgens et al 1967). Similarly, stimulation of the BNST elicits calls in the rhesus macaque (Robinson 1967) and squirrel monkey (Jurgens and Ploog 1970). In *Xenopus* frogs, stimulation of the CeA and BNST evokes fictive calling (Hall et al 2013).

### The cortex

We found three distinct cortical regions that were labelled with PRV – M1 motor cortex, prelimbic cortex (PrL), and entorhinal cortex (Ent) (Figure 12a-c). The “laryngeal motor cortex” (LMC), a region of M1 near the bregma, was infected only on the side contralateral to the injection, a finding consistent with studies of lab mice (Komiyama et al 2010, Arriaga et al 2012, and Chabout et al 2015). The rodent LMC is thought to be homologous to human LMC, which is essential for speech (Simonyan 2014). To our knowledge, our study is the first to examine whether the M1 neurons that connect to laryngeal muscles are also connected to other muscles important for vocalization. Because most M1 neurons were double-labelled, the region might better be regarded as a vocal motor cortex, rather than laryngeal cortex *per se*. The rodent prelimbic cortex (PrL) has also recently been implicated in vocalization. Working in rats, Bennett et al. (2019) argue that the posterior prelimbic cortex is connected to the LPAG, and that it may be homologous to the monkey ACC, which has long been recognized as an important area for vocal output in primates (Jurgens 2009). In *S.teguina,* we find that specific regions within the PrL show bilateral, co-infected neurons. This labeling, however, shows up sparsely at 96hpi, the final time point sampled. Lastly, a region of the entorhinal cortex (Ent) is infected bilaterally. Unlike the other regions, however, its label derives predominantly from the larynx. Although not canonically a “vocal” structure, it has been previously reported as being positively infected by a larynx-infected PRV circuit in the brain of rats (Van Daele and Cassell 2009).

### Uses and limitations of PRV

The primary advantage of our dual-PRV approach is the rapid, circuit-level delineation of brain regions and neurons that modulate one or both muscles (Callaway 2008, Hogue et al 2018, Saleeba et al 2019). We can also ask whether the timing of infection is consistent with networks identified in other species. Before discussing such interpretations, it is worthwhile to provide some caveats. The first is that in dual-labeling studies with virally infected neurons, the presence of one virus infection might limit infection by a second, a phenomenon known as the “principle of exclusion” or “superinfection inhibition” (Kobiler et al 2010, Doslikova et al 2019, Saleeba et al 2019). The widespread and region-specific presence of double-labelling in our study, however, suggests this may not be a major concern (Figure 13). A second caveat is that there is evidence that the virus may sometimes infect fibers of passage (Chen et al 1999). However, this finding stemmed from local injection of PRV into a brain region. In our study, targeting of muscles makes this form of infection unlikely. A third caveat is that PRV is large viral particle and is thought to be heavily biased toward synapses that are on or near the soma. This would not invalidate our infected regions but would suggest that they are a subset of the total vocal circuit. Finally, the time of arrival of the virus provides only a rough estimate of connectivity. While this approach is thus poorly suited for definitively characterizing projections, it is an excellent way to quickly delineate a circuit in a novel species and compare results to existing networks.

Although we targeted the cricothyroid and digastricus muscles, our results are remarkable concordant with a detailed single-label anatomical study that injected PRV into another intrinsic laryngeal muscle, the thyroarytenoid (Van Daele and Cassell 2009). Although the study was done in rats, both the identified brain regions and their times of arrival were quite similar to those we report. One notable difference is that these authors report ipsilateral infection for several forebrain structures – we found no strictly ipsilateral connections above the level of the brainstem, though most *Scotinomys* forebrain structures labelled more strongly ipsilaterally than contralaterally. Both studies, however, show some bilateral infection of most forebrain structures, so the overall pattern is one that is bilateral but biased toward ipsilateral infection. The uniquely contralateral infection in M1 cortex in singing mice and lab rats is one exception to this overall trend (Van Daele and Cassell 2009; see also Arriaga et al 2012 for similar results with lab mice).

### Conclusions

Taken together, our findings outline a vocal motor circuit that spans from motor neurons to limbic cortex. At the level of the hypothalamus, amygdala, midbrain, brainstem and spinal cord, the regions infected by the virus are remarkably concordant with vocal circuits reported for a variety of vertebrates. The CeA, MPOA, and BNST, for example, all regulate vocalization in frogs (Remage Healey and Bass 2008). Similarly, the PAG is a recurring and central player in the regulation of vocalization across vertebrate species (Kittleberger 2006, 2013). Our identification of cortical structures is consistent with recent reports in laboratory rodents and established circuits of vocalization in primates (Bennett et al. 2019, Jurgens 2009). Such patterns suggest that these cortical contributions likely date to at least the common ancestor of the clade euarchontoglires – the mammalian group defined to include rodents, tree shrews, bushbabies and primates – though they may prove to be older still. It would be particularly interesting to know how ancient these cortical/pallial circuits are. Outside of songbirds, whose vocal circuits are derived and specialized (Sakata and Yazaki-Sugiyama 2020), little work has been done to characterize pallial components of vocal circuits in non-mammals. These would clearly be interesting areas for further research. Overall our data, and the literature as a whole, suggests a general conservation of vocal circuitry across broad taxonomic scales.

Given the elaboration of vocalization in singing mice, and the fact that they are separated from laboratory rodents by roughly 40 million years of evolution (Steppan and Schenk 2017), we thought that we might see novel aspects of vocal circuitry. In contrast, the architecture of vocalization seems broadly similar to that reported in laboratory rodents. Thus the elaboration of vocalization does not seem to have been accompanied by gross changes in neural architecture – or at least none discernible with our current methods. We find three possible exceptions. One is the large doubly-labelled field within the reticular formation (LPGi, Gi, ROb), which we hypothesize may play an important role in the patterning of *Scotinomys* song. This seems likely to be a novel elaboration of vocal circuitry in the singing mouse. A second is the high level of bilateral infection we see in this circuit compared to some other reports (Doslikova et al 2019, Hettigoda et al 2015). The third and final difference is that most models of mammalian vocalization posit that descending forebrain projections from the hypothalamus influence vocalization through effects on the PAG (Jurgens REF, Tschida REF); in our data, however, robust double-labeling within PAG appeared either coincident with or following double-labeling in the hypothalamus – a pattern that suggests the hypothalamus may have some modulatory role that is not mediated by the PAG. To discern whether these anatomical features are indeed unique to *Scotinomys* will require more systematic study across relevant taxa.

In addition to improving our understanding of the evolution and conservation of mammalian and vertebrate vocal circuits, we also hoped to gain insights into the neural mechanisms that might underlie the complex decisions that mediate the adaptive modulation of advertisement displays. Among singing mice, as in many other taxa, display effort is influenced by reproductive state and body condition (Burkhard et al 2018, Pasch et al 2011, Giglio and Phelps 2020). In another paper (Zheng et al. 2021), we identify the presence of androgen receptors across this broad circuit, providing a potential mechanism for the androgen-dependence of *Scotinomys* song (Pasch et al. 2011). We have also found that body condition in general, and circulating leptin specifically, is associated with individual differences vocal effort. In this context, the extensive double labeling in the PVN is particularly interesting. The structure is at the nexus of limbic regions known as the social behavior network (Newman 1999), and the hypothalamic circuits of energy balance (Morton 2006). This structure seems to be a good candidate for the integration of social, reproductive and energetic information related to display in general, and to vocal behavior in particular.

In summary, this study employed a combination of viruses to target two distinct muscles in the larynx and jaw and identify neurons that were double-labelled by both retrograde viral tracers. Although known motor neurons were singly labelled, as expected, we found extensive double labelling at many higher levels. This circuit tracing suggests novel candidates for CPGs driving song, common limbic structures identified in other vertebrate groups, and cortical structures recently implicated in the control of vocalization in laboratory rodents as well as primates. We find that the circuitry seems broadly conserved across species and find no evidence of major circuit rearrangements associated with the elaboration of song in this novel model species. Lastly, our anatomical data suggest candidate brain regions for the integration of interoceptive and exteroceptive cues needed to produce adaptive variation in vocal behavior. We suggest that the dual-label approach, with PRV as well as with other transynaptic tracers, is an excellent way to quickly characterize vocal circuits in non-model mammals. Such studies will be important to our understanding of natural diversity in brain, behavior and evolution.

## Acknowledgements

We would like to thank Dr. Lynn Enquist for generously providing PRV-152 and PRV-614 through the CNNV. We would also like to thank Dr. Gustavo Arriaga for consultation on the cricothyroid injection. Funding for this project was made possible through P40 OD010996 to CNNV, 5R01NS113071-02 to MAL and SMP, NSF IOS-1457350 to SMP.

## Reference

1. Adkins-Regan, E. (2005). Hormones and Animal Social Behavior (1st ed.). Princeton: Princeton University Press.

2. Arriaga, G. (2012). Of Mice, Birds, and Men: The Mouse Ultrasonic Song System and Vocal Behavior. *Dissertation Abstracts International, B: Sciences and Engineering*, *72*(7), 3859. https://doi.org/10.1371/journal.pone.0046610

3. Arriaga, G., Macopson, J. J., & Jarvis, E. D. (2015). trans-synaptic Tracing from Peripheral Targets with Pseudorabies Virus Followed by Cholera Toxin and Biotinylated Dextran Amines Double Labeling 2. Surgical Preparation for Injections into Muscle. (September), 1–9. https://doi.org/10.3791/50672

4. Aston-Jones, G., & Cohen, J. D. (2005). AN INTEGRATIVE THEORY OF LOCUS COERULEUS-NOREPINEPHRINE FUNCTION: Adaptive Gain and Optimal Performance. Annual Review of Neuroscience, 28(1), 403–450. https://doi.org/10.1146/annurev.neuro.28.061604.135709

5. Bandler, R., & Keay, K. A. (1996). Columnar organization in the midbrain periaqueductal gray and the integration of emotional expression. Progress in Brain Research, 107, 285–300. https://doi.org/10.1016/s0079-6123(08)61871-3

6. Banerjee, A., Phelps, S. M., & Long, M. A. (2019). Singing mice. Current Biology, 29(6), R190–R191. https://doi.org/10.1016/j.cub.2018.11.048

7. Banfield, B. W., Kaufman, J. D., Randall, J. A., & Pickard, G. E. (2003). Development of pseudorabies virus strains expressing red fluorescent proteins: new tools for multisynaptic labeling applications. Journal of Virology, 77(18), 10106–10112. https://doi.org/10.1128/jvi.77.18.10106-10112.2003

8. Barkan, C. L., & Zornik, E. (2020). Inspiring song: The role of respiratory circuitry in the evolution of vertebrate vocal behavior. Developmental Neurobiology, 80(1–2), 31–41. https://doi.org/10.1002/dneu.22752

9. Barlow, S. M. (2009). Central pattern generation involved in oral and respiratory control for feeding in the term infant. Current Opinion in Otolaryngology & Head and Neck Surgery, 17(3), 187–193. https://doi.org/10.1097/MOO.0b013e32832b312a

10. Barrett, R. T., Bao, X., Miselis, R., & Altschuler, S. (1994). Brain Stem Localization of Rodent Esophageal Premotor. Gastroenterology, 107(3), 728–737.

11. Bass, A. H. (2014). ScienceDirect Central pattern generator for vocalization : is there a vertebrate morphotype ? Current Opinion in Neurobiology, 28, 94–100. https://doi.org/10.1016/j.conb.2014.06.012

12. Bass, A. H., & Remage-healey, L. (2008). Central pattern generators for social vocalization : Androgen-dependent neurophysiological mechanisms. 53, 659–672. https://doi.org/10.1016/j.yhbeh.2007.12.010

13. Bennett, P. J. G., Maier, E., & Brecht, M. (2019). Involvement of rat posterior prelimbic and cingulate area 2 in vocalization control. European Journal of Neuroscience, 50(7), 3164–3180. https://doi.org/10.1111/ejn.14477

14. Brainard, M. S., & Doupe, A. J. (2013). Translating Birdsong: Songbirds as a Model for Basic and Applied Medical Research. Annual Review of Neuroscience, 36(1), 489–517. https://doi.org/10.1146/annurev-neuro-060909-152826

15. Burkhard, T. T., Westwick, R. R., & Phelps, S. M. (2018). Adiposity signals predict vocal effort in Alston’s singing mice. Proceedings of the Royal Society B: Biological Sciences, 285(1877). https://doi.org/10.1098/rspb.2018.0090

16. Campbell, P., Pasch, B., Pino, J. L., Crino, O. L., Phillips, M., & Phelps, S. M. (2010). Geographic variation in the songs of neotropical singing mice: Testing the relative importance of drift and local adaptation. Evolution, 64(7), 1955–1972. https://doi.org/10.1111/j.1558-5646.2010.00962.x

17. Card, J. P., & Enquist, L. W. (2014). Transneuronal circuit analysis with pseudorabies viruses. Current Protocols in Neuroscience, 68, 1.5.1-1.5.39. https://doi.org/10.1002/0471142301.ns0105s68

18. Cassell, M. D. (2010). NIH Public Access. 162(2), 501–524. https://doi.org/10.1016/j.neuroscience.2009.05.005.Multiple

19. Chen, Z., & Wiens, J. J. (2020). The origins of acoustic communication in vertebrates. Nature Communications, 11(1), 1–8. https://doi.org/10.1038/s41467-020-14356-3

20. Connell, L. A. O., & Hofmann, H. A. (2011). The Vertebrate Mesolimbic Reward System and Social Behavior Network : A Comparative Synthesis. 3639, 3599–3639. https://doi.org/10.1002/cne.22735

21. Crino, O. L., Larkin, I., & Phelps, S. M. (2010). Stress coping styles and singing behavior in the short-tailed singing mouse (Scotinomys teguina). Hormones and Behavior, 58(2), 334–340. https://doi.org/10.1016/j.yhbeh.2010.02.011

22. Cummings, M. E., & Endler, J. A. (2018). 25 Years of sensory drive: The evidence and its watery bias. Current Zoology, 64(4), 471–484. https://doi.org/10.1093/cz/zoy043

23. Doslikova, B., Tchir, D., McKinty, A., Zhu, X., Marks, D. L., Baracos, V. E., & Colmers, W. F. (2019). Convergent neuronal projections from paraventricular nucleus, parabrachial nucleus, and brainstem onto gastrocnemius muscle, white and brown adipose tissue in male rats. Journal of Comparative Neurology, 527(17), 2826–2842. https://doi.org/10.1002/cne.24710

24. Dujardin, E., & Jürgens, U. (2006). Call type-specific differences in vocalization-related afferents to the periaqueductal gray of squirrel monkeys (Saimiri sciureus). Behavioural Brain Research, 168(1), 23–36. https://doi.org/10.1016/j.bbr.2005.10.006

25. Esposito, A., Demeurisse, G., Alberti, B., & Fabbro, F. (1999). Complete mutism after midbrain periaqueductal gray lesion. Neuroreport, 10(4), 681–685. https://doi.org/10.1097/00001756-199903170-00004

26. Falkner, A. L., Wei, D., Song, A., Watsek, L. W., Chen, I., Chen, P., … Lin, D. (2020). Hierarchical Representations of Aggression in a Hypothalamic-Midbrain Circuit. Neuron, 106(4), 637–648.e6. https://doi.org/10.1016/j.neuron.2020.02.014

27. Farley, G. R., Barlow, S. M., & Netsell, R. (1992). Factors influencing neural activity in parabrachial regions during cat vocalizations. Experimental Brain Research, 89(2), 341–351. https://doi.org/10.1007/BF00228250

28. Fay, R. A., & Norgren, R. (1997). *Identification of rat brainstem multisynaptic connections to the oral motor nuclei in the rat using pseudorabies virus II. Facial muscle motor systems*.

29. Fernández-Vargas, M., Tang-Martínez, Z., & Phelps, S. M. (2011). Singing, allogrooming, and allomarking behaviour during inter- and intra-sexual encounters in the Neotropical short-tailed singing mouse (Scotinomys teguina). Behaviour, 148(8), 945–965. https://doi.org/10.1163/000579511X584591

30. Foote, S. L., Freedman, R., & Oliver, A. P. (1975). Effects of putative neurotransmitters on neuronal activity in monkey auditory cortex. Brain Research, 86(2), 229–242. https://doi.org/10.1016/0006-8993(75)90699-x

31. Franklin, K. B.., & Paxinos, G. (2007). The Mouse Atlas In Stereotaxic Coordinates (3rd ed.). New York.

32. Fusani, L., Barske, J., Day, L. D., Fuxjager, M. J., & Schlinger, B. A. (2014). Physiological control of elaborate male courtship: Female choice for neuromuscular systems. Neuroscience and Biobehavioral Reviews, 46(P4), 534–546. https://doi.org/10.1016/j.neubiorev.2014.07.017

33. Gao, S.-C., Wei, Y.-C., Wang, S.-R., & Xu, X.-H. (2019). Medial Preoptic Area Modulates Courtship Ultrasonic Vocalization in Adult Male Mice. Neuroscience Bulletin, 35(4), 697–708. https://doi.org/10.1007/s12264-019-00365-w

34. Gentner, T. Q., & Margoliash, D. (2006). The Neuroethology of Vocal Communication: Perception and Cognition. Acoustic Communication, 324–386. https://doi.org/10.1007/0-387-22762-8_7

35. Goodson, J. L. (2005). The vertebrate social behavior network : Evolutionary themes and variations i. 48, 11–22. https://doi.org/10.1016/j.yhbeh.2005.02.003

36. Goodson, J. L., & Bass, A. H. (2000). Forebrain peptides modulate sexually polymorphic vocal circuitry. Nature, 403(6771), 769–772. https://doi.org/10.1038/35001581

37. Goodson, J. L., & Bass, A. H. (2002). Vocal-acoustic circuitry and descending vocal pathways in teleost fish: Convergence with terrestrial vertebrates reveals conserved traits. Journal of Comparative Neurology, 448(3), 298–322. https://doi.org/10.1002/cne.10258

38. Gruber-Dujardin, E. (2010). Role of the periaqueductal gray in expressing vocalization. In S. M. Brudzynski (Ed.), Handbook of Mammalian Vocalization (1st ed., pp. 313–328). London: Elsevier.

39. Guyenet, P. G., Koshiya, N., Huangfu, D., Verberne, A. J. M., & Riley, T. A. (1993). Central respiratory control of A5 and A6 pontine noradrenergic neurons. American Journal of Physiology - Regulatory Integrative and Comparative Physiology, *264*(6 33-6). https://doi.org/10.1152/ajpregu.1993.264.6.r1035

40. Hage, S. R. (2010). Localization of the central pattern generator for vocalization. In S. M. Brudzynski (Ed.), Handbook of Mammalian Vocalization (1st ed., pp. 329–337). London: Elsevier.

41. Hage, S. R., & Nieder, A. (2016). Dual Neural Network Model for the Evolution of Speech and Language. Trends in Neurosciences, 39(12), 813–829. https://doi.org/10.1016/j.tins.2016.10.006

42. Hall, I. C., Ballagh, I. H., & Kelley, D. B. (2013). The Xenopus amygdala mediates socially appropriate vocal communication signals. Journal of Neuroscience, 33(36), 14534–14548. https://doi.org/10.1523/JNEUROSCI.1190-13.2013

43. Hamed, A., & Boguszewski, P. M. (2018). Effects of Morphine and Other Opioid Ligands on Emission of Ultrasonic Vocalizations in Rats. In Handbook of Behavioral Neuroscience (1st ed., Vol. 25). https://doi.org/10.1016/B978-0-12-809600-0.00031-7

44. Herbert, H., Moga, M. M., & Saper, C. B. (1990). Connections of the parabrachial nucleus with the nucleus of the solitary tract and the medullary reticular formation in the rat. The Journal of Comparative Neurology, 293(4), 540–580. https://doi.org/10.1002/cne.902930404

45. Hernandez-Miranda, L. R., Ruffault, P. L., Bouvier, J. C., Murray, A. J., Morin-Surun, M. P., Zampieri, N., … Birchmeier, C. (2017). Genetic identification of a hindbrain nucleus essential for innate vocalization. Proceedings of the National Academy of Sciences of the United States of America, 114(30), 8095–8100. https://doi.org/10.1073/pnas.1702893114

46. Hettigoda, N. S., Fong, A. Y., Badoer, E., McKinley, M. J., Oldfield, B. J., & Allen, A. M. (2015). Identification of CNS neurons with polysynaptic connections to both the sympathetic and parasympathetic innervation of the submandibular gland. Brain Structure and Function, 220(4), 2103–2120. https://doi.org/10.1007/s00429-014-0781-1

47. Hilaire, G., Viemari, J.-C., Coulon, P., Simonneau, M., & Bévengut, M. (2004). Modulation of the respiratory rhythm generator by the pontine noradrenergic A5 and A6 groups in rodents. Respiratory Physiology & Neurobiology, 143(2–3), 187–197. https://doi.org/10.1016/j.resp.2004.04.016

48. Hisa, Y. (2016). Neuroanatomy and neurophysiology of the larynx. Neuroanatomy and Neurophysiology of the Larynx, 1–123. https://doi.org/10.1007/978-4-431-55750-0

49. Hisa, Y., Sato, F., Fukui, K., Ibata, Y., & Mizuokoshi, O. (1984). Nucleus Ambiguus Motoneurons Innervating the Canine Intrinsic Laryngeal Muscles by the Fluorescent Labeling Technique. Experimental Neurology, 84, 441–449.

50. Hogue, I. B., Card, J. P., Rinaman, L., Staniszewska Goraczniak, H., & Enquist, L. W. (2018). Characterization of the neuroinvasive profile of a pseudorabies virus recombinant expressing the mTurquoise2 reporter in single and multiple injection experiments. Journal of Neuroscience Methods, 308(August), 228–239. https://doi.org/10.1016/j.jneumeth.2018.08.004

51. Hooper, E. T., & Carleton, M. D. M. (1976). Reproduction, growth and development in two contiguously allopatric rodent species, genus Scotinomys. *Miscellaneous Publications, Museum of Zoology*, University of Michigan, 151(15), 1–52. Retrieved from http://deepblue.lib.umich.edu/handle/2027.42/56395

52. Iv, E. S., Cheng, S., Takatoh, J., Han, B., & Wang, F. (2014). Monosynaptic premotor circuit tracing reveals neural substrates for oro-motor coordination. 1–23. https://doi.org/10.7554/eLife.02511

53. Jovanovic, K., Pastor, A. M., & O’Donovan, M. J. (2010). The use of PRV-bartha to define premotor inputs to lumbar motoneurons in the neonatal spinal cord of the mouse. PLoS ONE, 5(7). https://doi.org/10.1371/journal.pone.0011743

54. Ju, U. (n.d.). *The Neural Control of Vocalization in Mammals : A Review*. https://doi.org/10.1016/j.jvoice.2007.07.005

55. Jürgens, U. (1994). The role of the periaqueductal grey in vocal behaviour. Behavioural Brain Research, 62(2), 107–117. https://doi.org/10.1016/0166-4328(94)90017-5

56. Jürgens, Uwe. (2002). Neural pathways underlying vocal control. Neuroscience and Biobehavioral Reviews, 26(2), 235–258. https://doi.org/10.1016/s0149-7634(01)00068-9

57. Kang, T. C., Lee, B. H., Seo, J., Song, S. H., Kim, J. S., Won, M. H., … Lee, H. S. (1999). The nuclei innervating digastric muscle do not project to the hypoglossal nucleus in the rat. *Anatomia, Histologia*, Embryologia, 28(1), 39–40. https://doi.org/10.1046/j.1439-0264.1999.00159.x

58. Kelley, D. B., Ballagh, I. H., Barkan, C. L., Bendesky, A., Elliott, T. M., Evans, B. J., … Zornik, E. (2020). Generation, coordination, and evolution of neural circuits for vocal communication. Journal of Neuroscience, 40(1), 22–36. https://doi.org/10.1523/JNEUROSCI.0736-19.2019

59. Kemplay, S., & Cavanagh, J. B. (1983). Bilateral innervation of the anterior digastric muscle by trigeminal motor neurons. Journal of Anatomy, 136(Pt 2), 417–423. Retrieved from https://pubmed.ncbi.nlm.nih.gov/6853354

60. Kingsbury, M. A., Kelly, A. M., Schrock, S. E., & Goodson, J. L. (2011). Mammal-Like Organization of the Avian Midbrain Central Gray and a Reappraisal of the Intercollicular Nucleus. 6(6). https://doi.org/10.1371/journal.pone.0020720

61. Koblinger, K., Füzesi, T., Ejdrygiewicz, J., Krajacic, A., Bains, J. S., & Whelan, P. J. (2014). Characterization of A11 neurons projecting to the spinal cord of mice. PloS One, 9(10), e109636. https://doi.org/10.1371/journal.pone.0109636

62. Lein, E. S., Hawrylycz, M. J., Ao, N., Ayres, M., Bensinger, A., Bernard, A., … Jones, A. R. (2007). Genome-wide atlas of gene expression in the adult mouse brain. Nature, 445(7124), 168–176. https://doi.org/10.1038/nature05453

63. Martins, A. R. O., & Froemke, R. C. (2015). Coordinated forms of noradrenergic plasticity in the locus coeruleus and primary auditory cortex. Nature Neuroscience, 18(10), 1483–1492. https://doi.org/10.1038/nn.4090

64. Miller, J. R., & Engstrom, M. D. (2007). Vocal Stereotypy and Singing Behavior in Baiomyine Mice. Journal of Mammalogy. https://doi.org/10.1644/06-MAMM-A-386R.1

65. Moore, J. D., Kleinfeld, D., & Wang, F. (2014). How the brainstem controls orofacial behaviors comprised of rhythmic actions. Trends in Neurosciences, 37(7), 370–380. https://doi.org/10.1016/j.tins.2014.05.001

66. Nevue, A. A., Felix 2nd, R. A., & Portfors, C. V. (2016). Dopaminergic projections of the subparafascicular thalamic nucleus to the auditory brainstem. Hearing Research, 341, 202–209. https://doi.org/10.1016/j.heares.2016.09.001

67. Newman, S. W. (n.d.). *The Medial Extended Amygdala in Male Reproductive Behavior A Node in the Mammalian Social Behavior Network*. 242–257.

68. Nieder, A., & Mooney, R. (2020). The neurobiology of innate, volitional and learned vocalizations in mammals and birds. Philosophical Transactions of the Royal Society B: Biological Sciences, 375(1789). https://doi.org/10.1098/rstb.2019.0054

69. Núñez-Abades, P. A., Portillo, F., & Pásaro, R. (1990). Characterisation of afferent projections to the nucleus ambiguus of the rat by means of fluorescent double labelling. Journal of Anatomy, 172, 1–15.

70. Okobi, D. E., Banerjee, A., Matheson, A. M. M., Phelps, S. M., & Long, M. A. (2019). Motor cortical control of vocal interaction in neotropical singing mice. Science, 363(6430), 983–988. https://doi.org/10.1126/science.aau9480

71. Okobi, D. E., & PhD. (2016). *A cortical locus modulates vocal motor sequences in Alston’s singing mouse (Scotinomys teguina) by Daniel Ebele Okobi, Jr. A dissertation submitted in partial fulfillment of the requirements for the degree of Doctor of Philosophy Department of Basic*.

72. Pasch, B., Bolker, B. M., & Phelps, S. M. (2013). Interspecific dominance via vocal interactions mediates altitudinal zonation in neotropical singing mice. The American Naturalist, 182(5), E161–73. https://doi.org/10.1086/673263

73. Pasch, B., George, A. S., Campbell, P., & Phelps, S. M. (2011a). Androgen-dependent male vocal performance in fl uences female preference in Neotropical singing mice. Animal Behaviour, 82(2), 177–183. https://doi.org/10.1016/j.anbehav.2011.04.018

74. Pasch, B., George, A. S., Campbell, P., & Phelps, S. M. (2011b). Androgen-dependent male vocal performance influences female preference in Neotropical singing mice. Animal Behaviour, 82(2), 177–183. https://doi.org/10.1016/j.anbehav.2011.04.018

75. Pérez, C. A., Stanley, S. A., Wysocki, R. W., Havranova, J., Ahrens-Nicklas, R., Onyimba, F., & Friedman, J. M. (2011). Molecular annotation of integrative feeding neural circuits. Cell Metabolism, 13(2), 222–232. https://doi.org/10.1016/j.cmet.2010.12.013

76. Prins, G. S., Birch, L., & Greene, G. L. (1991). Androgen receptor localization in different cell types of the adult rat prostate. Endocrinology, 129(6), 3187–3199. https://doi.org/10.1210/endo-129-6-3187

77. Remage-Healey, L., Maidment, N. T., & Schlinger, B. A. (2008). Forebrain steroid levels fluctuate rapidly during social interactions. Nature Neuroscience, 11(11), 1327–1334. https://doi.org/10.1038/nn.2200

78. Rhodes, H. J., Yu, H. J., & Yamaguchi, A. (2007). Xenopus vocalizations are controlled by a sexually differentiated hindbrain central pattern generator. The Journal of Neuroscience : The Official Journal of the Society for Neuroscience, 27(6), 1485–1497. https://doi.org/10.1523/JNEUROSCI.4720-06.2007

79. Riede, T. (2013). Stereotypic laryngeal and respiratory motor patterns generate different call types in rat ultrasound vocalization. *Journal of Experimental Zoology. Part A*, Ecological Genetics and Physiology, 319(4), 213–224. https://doi.org/10.1002/jez.1785

80. Riede, T., & Pasch, B. (2020). Pygmy mouse songs reveal anatomical innovations underlying acoustic signal elaboration in rodents. The Journal of Experimental Biology, 223(12), jeb223925. https://doi.org/10.1242/jeb.223925

81. Samuels, E. R., & Szabadi, E. (2008). Functional neuroanatomy of the noradrenergic locus coeruleus: its roles in the regulation of arousal and autonomic function part I: principles of functional organisation. Current Neuropharmacology, 6(3), 235–253. https://doi.org/10.2174/157015908785777229

82. Saper, C. B., & Loewy, A. D. (1980). Efferent connections of the parabrachial nucleus in the rat. Brain Research, 197(2), 291–317. https://doi.org/10.1016/0006-8993(80)91117-8

83. Sara, S. J. (2009). The locus coeruleus and noradrenergic modulation of cognition. Nature Reviews Neuroscience, 10(3), 211–223. https://doi.org/10.1038/nrn2573

84. Saravanan, V., Hoffmann, L. A., Jacob, A. L., Berman, G. J., & Sober, S. J. (2019). Dopamine depletion affects vocal acoustics and disrupts sensorimotor adaptation in songbirds. ENeuro, 6(3). https://doi.org/10.1523/ENEURO.0190-19.2019

85. Schlinger, B. A., Paul, K., & Monks, D. A. (2018). Muscle, a conduit to brain for hormonal control of behavior. Hormones and Behavior, 105(August), 58–65. https://doi.org/10.1016/j.yhbeh.2018.07.002

86. Schmidt, R. S. (1968). Preoptic Activation of Frog Mating Behavior. Behaviour, 30(2/3), 239–257. Retrieved from http://www.jstor.org/stable/4533213

87. Schwartz, C. P., & Smotherman, M. S. (2011). Mapping vocalization-related immediate early gene expression in echolocating bats. Behavioural Brain Research, 224(2), 358–368. https://doi.org/10.1016/j.bbr.2011.06.023

88. Simonyan, K., Horwitz, B., & Jarvis, E. D. (2012). Dopamine regulation of human speech and bird song: a critical review. Brain and Language, 122(3), 142–150. https://doi.org/10.1016/j.bandl.2011.12.009

89. Smotherman, M., Schwartz, C., & Metzner, W. (2010). Vocal-respiratory interactions in the parabrachial nucleus. In S. M. Brudzynski (Ed.), Handbook of Mammalian Vocalization (1st ed., pp. 383–392). London: Elsevier.

90. Stanley, S., Pinto, S., Segal, J., Pérez, C. A., Viale, A., DeFalco, J., … Friedman, J. M. (2010). Identification of neuronal subpopulations that project from hypothalamus to both liver and adipose tissue polysynaptically. Proceedings of the National Academy of Sciences of the United States of America, 107(15), 7024–7029. https://doi.org/10.1073/pnas.1002790107

91. Suthers, R. A., Fitch, W. T., Fay, R. R., & Popper, A. N. (2004). Vertebrate sound production and acoustic communication. In (Fay RR & Popper AN (ed) Springer Handbook of Auditory Research. The Senses of Fish, 19–49. https://doi.org/10.1007/978-3-319-27721-9

92. Szabadi, E. (2013). Functional neuroanatomy of the central noradrenergic system. Journal of Psychopharmacology, 27(8), 659–693. https://doi.org/10.1177/0269881113490326

93. Tobiansky, D. J., & Fuxjager, M. J. (2020). Sex Steroids as Regulators of Gestural Communication. Endocrinology, 161(7), 1–12. https://doi.org/10.1210/endocr/bqaa064

94. Tschida, K., Michael, V., Takatoh, J., Han, B. X., Zhao, S., Sakurai, K., … Wang, F. (2019). A Specialized Neural Circuit Gates Social Vocalizations in the Mouse. Neuron, 103(3), 459–472.e4. https://doi.org/10.1016/j.neuron.2019.05.025

95. Waldbaum, S., Hadziefendic, S., Erokwu, B., Zaidi, S. I. A., & Haxhiu, M. A. (2001). CNS innervation of posterior cricoarytenoid muscles: A transneuronal labeling study. Respiration Physiology, 126(2), 113–125. https://doi.org/10.1016/S0034-5687(01)00200-6

96. Wee, N. K. Y., Lorenz, M. R., Bekirov, Y., Jacquin, M. F., & Scheller, E. L. (2019). Shared autonomic pathways connect bone marrow and peripheral adipose tissues across the central neuraxis. Frontiers in Endocrinology, 10(SEP), 1–16. https://doi.org/10.3389/fendo.2019.00668

97. Wei, Y. C., Wang, S. R., Jiao, Z. L., Zhang, W., Lin, J. K., Li, X. Y., … Xu, X. H. (2018). Medial preoptic area in mice is capable of mediating sexually dimorphic behaviors regardless of gender. Nature Communications, 9(1). https://doi.org/10.1038/s41467-017-02648-0

98. Wiedmann, N. M., Stefanidis, A., & Oldfield, B. J. (2017). Characterization of the central neural projections to brown, white, and beige adipose tissue. FASEB Journal, 31(11), 4879–4890. https://doi.org/10.1096/fj.201700433R

99. Zhang, Y. S., & Ghazanfar, A. A. (2020). A Hierarchy of Autonomous Systems for Vocal Production. Trends in Neurosciences, 43(2), 115–126. https://doi.org/10.1016/j.tins.2019.12.006

